# Age, Origin, and Biogeography: Unveiling the Factors Behind the Diversification of Dung Beetles

**DOI:** 10.1101/2021.01.26.428346

**Authors:** Orlando Schwery, Brian C. O’Meara

## Abstract

The remarkable diversity and global distribution of dung beetles has long attracted the interest of researchers. However, there is still an ongoing debate on their origin, the reasons behind their diversity, and their path to global distribution. The two most prominent hypotheses regarding their origin and biogeographic history involve either vicariance events after the breakup of Gondwana, or an African origin and subsequent dispersal. One of the key reasons why the question is still disputed is a dependence on knowing the age of the dung beetles – a Mesozoic origin would favor the scenario of Gondwanan vicariance, a Cenozoic origin would suggest the out-of-Africa scenario. To help settle this longstanding question, we provide a taxonomically expanded phylogeny, with divergence times estimated under two calibration schemes suggesting an older or younger origin respectively. Using model-based inference, we estimate the ancestral area of the group and test for the influence of ranges on diversification rates. Our results support the hypothesis of an old age for Scarabaeinae and Gondwanan origin but remain ambiguous about the exact relation of range on lineage diversification.

## Introduction

Dung beetles are a remarkable group of insects. Their unusual lifestyle requiring the dung of other animals to feed and reproduce gave rise to a host of morphological and behavioral specializations, as adaptations to the various ecological peculiarities they face in their worldwide distribution (Hanski & Cambefort 1991). While their diversity of around 5,300 species is comparatively modest among beetles, their dung-processing activity makes them one of the most important groups of insects, both ecologically (Nichols *et al*. 2008) and economically (Losey & Vaughan 2006). Their unusual life history has attracted considerable interest of researchers, in ecology and evolution (Hanski & Cambefort 1991; Scholtz, Davis & Kryger 2009), conservation (Spector 2006), and even developmental biology (Moczek 2011). Despite that, key aspects concerning their origin and the factors behind their diversity are still unknown and subject of debate in the field. The ecology of extant species has been well studied (Hanski & Cambefort 1991), but the main hindrance to understanding historical, evolutionary aspects of their biology is the lack of a comprehensive and reliable phylogeny (Tarasov & Génier 2015), and with that, validation for their taxonomy (Tarasov & Dimitrov 2016).

Probably the two most debated questions of dung beetle evolution revolve around their geographic origin and what led to their current diversity and distribution. A relation to their associated dung producers is suspected (Scholtz, Davis & Kryger 2009; Gunter *et al*. 2016), but disagreement exists over whether the success of dung beetles has always been an association with mammals, or whether dinosaurs were involved early in their history. Regarding their geographic origin and subsequent spread, the main competing hypotheses are whether it was Gondwanan-vicariance (Davis, Scholtz & Philips 2002) or dispersal out of Africa (Sole & Scholtz 2010). The answers to all these hypotheses partially hinge on the question of how old the group is. It has long been debated whether Scarabaeinae are of Mesozoic or Cenozoic origin – the former would make Gondwanan vicariance and feeding on dinosaur dung plausible, the latter would rule it out. Earlier attempts to determine the age of dung beetles using various approaches led to widely different estimates, ranging from mid Mesozoic to early Cenozoic (Hanski & Cambefort 1991; Scholtz & Chown 1995; Krell 2000; Davis, Scholtz & Philips 2002; Krell 2006). More recent attempts using phylogenies also ranged from Cretaceous to Eocene-Oligocene (Wirta, Orsini & Hanski 2008; Ahrens, Schwarzer & Vogler 2014; Mlambo, Sole & Scholtz 2015; Gunter *et al*. 2016), shifting the support for early and late origins of Scarabaeinae over time (Scholtz, Davis & Kryger 2009). At this time, the field is still divided: an old Mesozoic origin of dung beetles, and with that a biogeographical scenario of Gondwanan vicariance and subsequent dispersal has been supported by some studies (Davis, Scholtz & Philips 2002; Gunter *et al*. 2016; Gunter *et al*. 2018), whereas a young Cenozoic origin and therefore an out-of-Africa dispersal scenario has been supported by others (Monaghan *et al*. 2007; Sole & Scholtz 2010; Davis, Scholtz & Sole 2016).

To address these questions, we provide an extended phylogeny of Scarabaeinae, with divergence times estimates based on calibrations reflecting two different assumptions of their maximal age. We use this phylogeny and the inferred age of the group to address the question of their geographical origin using model-based methods for ancestral range estimation; and the question of how it relates to their lineage diversification using range-dependent diversification rate estimation. In particular, focusing on their Gondwanan origin and subsequent dispersals to areas of (what was formerly) Laurasia and Madagascar, we address the question of whether the dispersal into new areas was promoting dispersal, or whether their place of origin is the main source of diversification. Finally, we test a hypothesis that links dung beetle diversity to the dung producers present in different regions (Davis & Scholtz 2001; Scholtz, Davis & Kryger 2009). Specifically, linking beetle diversity to the size of mammal dung producers, and the diversity of different dung types they produced, suggesting that areas with larger mammals and more diverse droppings would allow for a more diverse dung beetle fauna.

## Materials and Methods

### Phylogenetic Analysis

We wrote a script in R (R Development Core Team 2014) using the packages *reutils* (Schöfl 2016) and *seqinr* (Charif *et al*. 2005) to query GenBank (Sayers *et al*. 2019) for any nucleotide sequences matching the organism label “Scarabaeinae” and that carried either “COI”, “16S”, “18S”, “28S”, or “CAD” in the title. We downloaded the resulting accessions using the packages *ape* (Paradis, Claude & Strimmer 2004) and *insect* (Wilkinson *et al*. 2018), removed any duplicates and saved them as FASTA files. The 28S accessions included both 28SD2 and 28SD3 sequences, which were sorted and aligned separately. The retrieved COI sequences largely covered two adjacent regions of the gene, and since only few accessions covered both regions, we decided to split them into two separate alignments (further called COI-1 and COI-2 respectively). If a marker had taxa with several accessions, the longest sequence was chosen unless it proved to be clearly different from the other accessions for that taxon. Taxa which were only determined to genus level were only included if they were the only representative of that genus.

The accessions of the seven markers were aligned separately using AliView v.1.26 (Larsson 2014): They were aligned using MAFFT globalpair, and then visually inspected and adjusted manually where necessary. During this process, any sequences that showed considerable mismatch with the rest of the alignment were submitted to BLAST (Altschul *et al*. 1997) to detect any mislabeled sequences from other organismal groups. Similarly, quick RAxML (Stamatakis 2014) runs were performed for each aligned marker separately, and were tested for generic monophyly and long branches. Generic monophyly was tested using the package *MonoPhy* (Schwery & O’Meara 2016). Long branches were determined using the package *ips*; terminal branches were considered exceptionally long if their length was more than 0.25 times the maximum tip height in the tree, or if the product of their length and height was more than two times the interquartile range away from the third quartile of all tip length and tip height products in the tree. Any taxa that stood out by branch length or placement were checked using BLAST as well. The accession numbers of the sequences used in the final alignment can be found in Table S1.

Poorly aligned or divergent regions in each alignment were removed using Gblocks (Castresana 2000) (settings: minimum bases for conserved regions >0.5x alignment length, minimum for flanking regions >0.7x alignment length, maximum contiguous nonconserved sites: 8, minimum block length: 4 (noncoding) or 5 (coding), gap positions allowed with > half of sequences having gap). Finally, all alignments were concatenated using the package *evobiR* (Blackmon & Adams 2015). Partitioning the alignment by each marker (the two marker parts in case of COI), we used PartitionFinder v.2.1.1 (Lanfear *et al*. 2017) to determine the best substitution models for each partition, using the greedy algorithm (Lanfear *et al*. 2012), and PhyML (Guindon *et al*. 2010). Additionally, the package *ClockstaR* (Duchene, Molak & Ho 2014) was used to determine whether the partitions should have different clock models.

Using the concatenated alignment and the determined substitution models, a maximum likelihood phylogeny was constructed with RAxML v.8.2.4 (Stamatakis 2014), using the rapid hill climbing algorithm with 100 rapid bootstrap samples. The ingroup and four clades that would later be used for fossil calibrations (see below) were constrained to be monophyletic. Because of branch support issues, we used the online implementation of RogueNaRok (Aberer, Krompass & Stamatakis 2013) to find potential rogue taxa. Taxa whose exclusion would lead to a raw improvement of more than 1 were inspected and 136 sequences were eventually dropped and a final RAxML tree was built from this reduced alignment. To account for weakly supported clade relationships in subsequent analyses, ten sets of bootstrapped alignments were created, for each of which a separate tree was built in the same manner as from the full alignment. Monophyly on genus and tribe level was assessed for all trees using the R package *MonoPhy* (Schwery & O’Meara 2016).

### Divergence Time Estimation

Reliable Scarabaeinae fossils that have recently been re-examined by Tarasov *et al*. (2016) were chosen to calibrate tree nodes during divergence time estimation. Of the 35 fossils previously assigned to Scarabaeinae, they considered only 21 to be assigned reliably on the basis of their morphological characters. From among these, the earliest fossils with confident generic placement were chosen for each genus that we could assign a node to. The minimal age of the fossils was used as minimum constraint for the stem age of the corresponding clade. The oldest reliable fossil dung beetle, *Lobateuchus parisii*, was used to constrain the subfamily, and the minimal age estimate for *Juraclopus rohendorfi*, the oldest fossil of the family Scarabaeidae, was used as the minimal age constraint for the crown of the whole tree.

In order to get times to constrain the maximal age of these clades, we consulted previous studies that had estimated the age of Scarabaeinae. On the younger end of the spectrum, some studies refer to Wirta, Orsini and Hanski (2008), or Mlambo, Sole and Scholtz (2015) for young estimates of the age of dung beetles (33.9ma or 56ma respectively). However, neither of those estimates are particularly useful to calibrate the age of the whole group, as Wirta, Orsini and Hanski (2008) focused on Helictopleurini of Madagascar, and Mlambo, Sole and Scholtz (2015) exclusively on African dung beetles. While some more dated phylogenies of clades within the subfamily exist, the only examples that include an age for the whole group are of higher taxa in which the dung beetles are nested. Ahrens, Schwarzer and Vogler (2014) constructed a tree of 146 taxa of Scarabaeoidea, in which the crown Scarabaeinae was estimated at 89.6ma (ranging from 83.5-98.1ma), while Gunter *et al*. (2016) in a phylogeny of 445 taxa of Scarabaeoidea estimated it to range from 118.8-131.6ma. While various studies estimated the ages within Scarabaeoidea (Ahrens, Schwarzer & Vogler 2014; McKenna *et al*. 2015; Toussaint *et al*. 2017), the ages of Scarabaeinae and Scarabaeidae could not always be obtained from them – consequently, we relied solely on Gunter *et al*. (2016) for an age estimate of Scarabaeidae (116.85-199.64ma).

Because no clade can be older than the clade in which it is nested, we constrained the maximal stem ages for all constrained genera (as well as the maximal crown age of all Scarabaeinae) to be the maximal estimated age of the Scarabaeinae, and the maximal crown age of the whole tree (being the stem age of Scarabaeinae) to be the maximal estimated age of Scarabaeidae. Given the disagreement of estimated ages in the literature, we set up two different sets of constraints: an ‘old’ one, with the Scarabaeinae and Scarabaeidae ages as estimated by Gunter *et al*. (2016), and a ‘young’ one, with the Scarabaeinae and Scarabaeidae ages as estimated by Ahrens, Schwarzer and Vogler (2014) – with the latter actually being the age of Scarabaeoidea, due to the lack of available ages for the actual Scarabaeidae (apart from that by Gunter *et al*. (2016)). While McKenna *et al*. (2015) actually estimated a younger age for Scarabaeoidea, their estimate is younger than the age of the oldest fossil of that group, which is why we chose to use the next older estimate by Ahrens, Schwarzer and Vogler (2014) ranging from 167.2-181.8ma. The different fossil constraints used can be seen in Table 2.

**Table 1.**
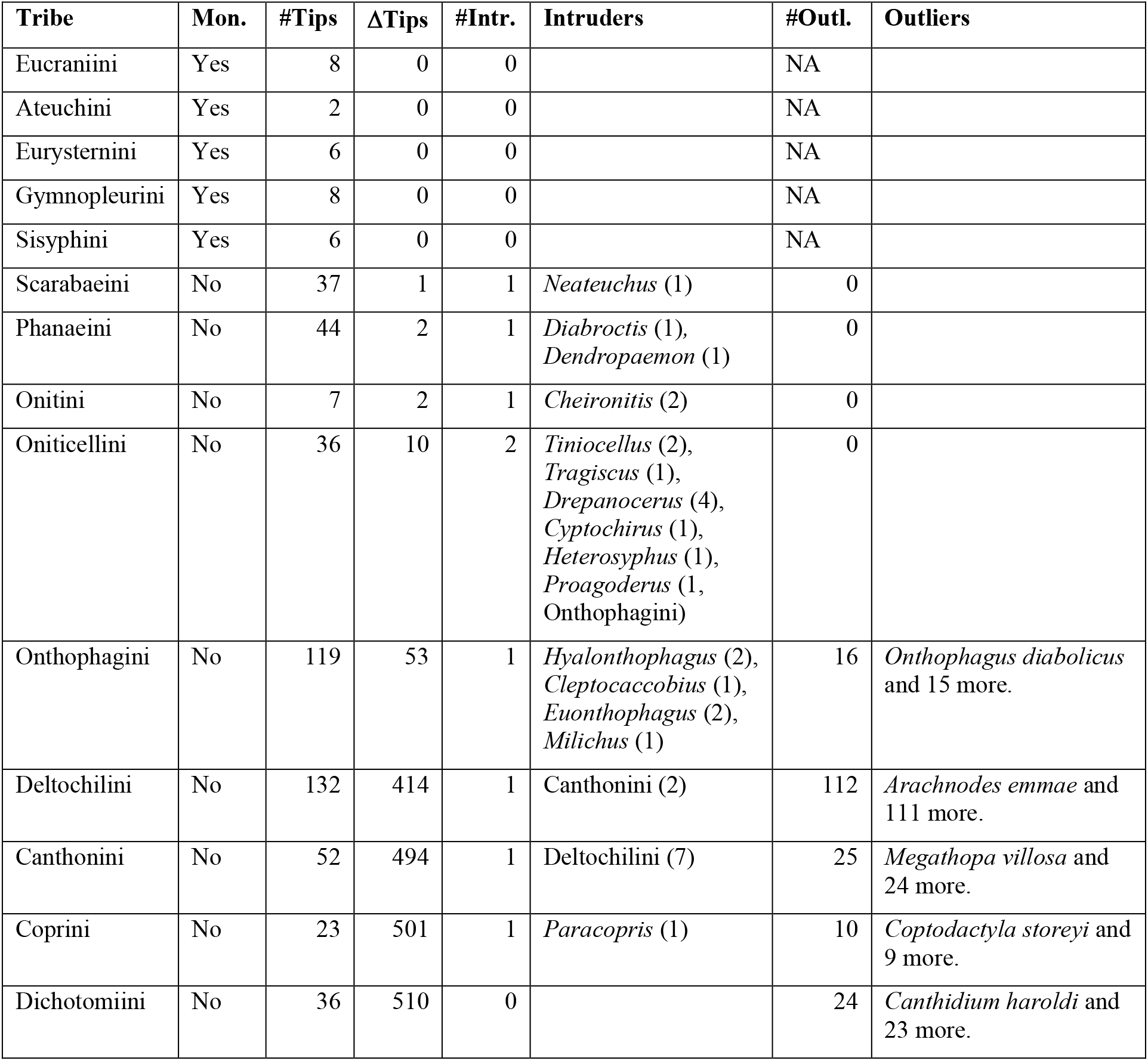
Tribal Monophyly-Issues Dung Beetles. Monophyly status and reasons for non-monophyly for all tribes in the full matrix trees. Mon.=monophyly-status, #Tips=number of taxa assigned to this tribe, ΔTips=number of additional taxa in this clade (descendants of same MRCA), #Intr=number of tribes among intruder taxa, Intruders=Genera or Tribes intruding (numbers in parentheses are numbers of tips, all taxa *incertae sedis* unless indicated otherwise), #Outl.=number of taxon-members outside the main clade.

**Table 2.**
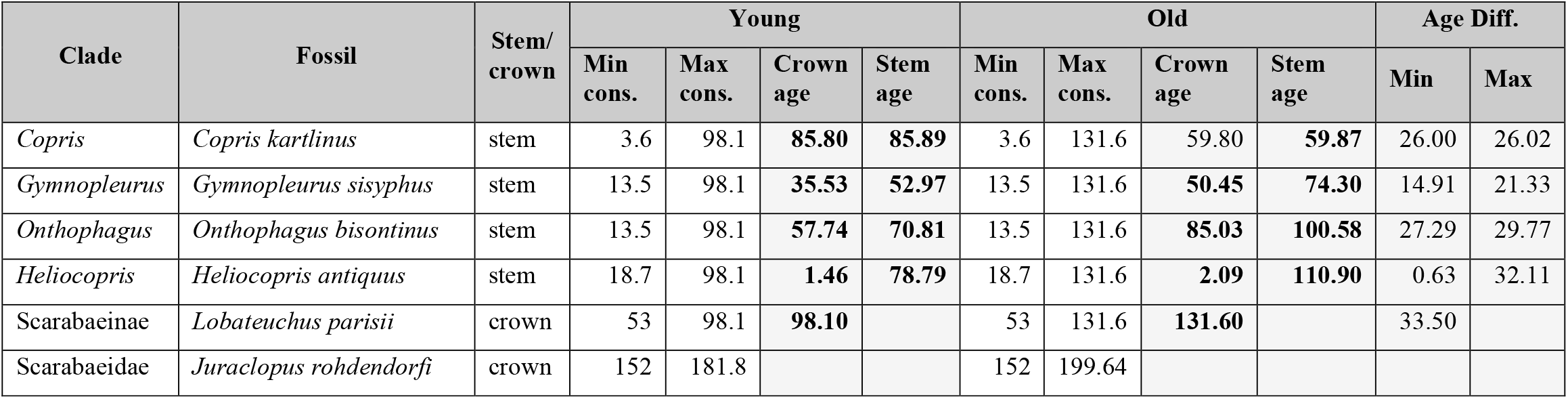
Node Calibrations and Estimated Ages. For each node age calibration, the corresponding clade, fossil, location on the branch (crown or stem) are given. For each calibration scheme (‘young’ vs. ‘old’), the age constraints used and the estimated node ages on crown and stem, are indicated for each calibrated node, followed by the difference between the estimates.

Those two sets of age constraints were then used to estimate divergence times for the ML tree obtained through RAxML and all ten bootstrap trees, using penalized likelihood in treePL (Smith & O’Meara 2012). We performed a thorough search with random cross validation, which was preceded by a preliminary run to estimate the best tuning parameters and smoothing factor.

### Occurrence Data and Ancestral Range Estimation

We used the package *rgbif* (Chamberlain *et al*. 2016; Chamberlain & Boettiger 2017) to download occurrence data from GBIF (http://data.gbif.org) for all taxa determined to species level. Using an R script, the retrieved records were cleaned from empty or invalid entries, with regards to the basis of record, identification date, country and coordinates, and sets with unique countries or coordinates per taxon respectively were made. Name validity of unavailable taxa was checked using the package *taxize* (Chamberlain & Szocs 2013), and eventually searched on genus level. For any taxa that were still missing, occurrence information was searched for in the literature.

We defined the areas taxa occur in as continents (Africa, Asia, Europe, North America, South America, Oceania), and defined Madagascar as a separate area, given the high number of taxa endemic to it (Miraldo, Wirta & Hanski 2011). However, we did not define India as a separate area, and considered it part of Asia. India has a peculiar tectonic history of initially staying connected to Madagascar until breaking off around 87ma and drifting northwards to collide with Asia around 55-43ma (Seton *et al*. 2012). Not explicitly including it as a separate area means a major simplification for the model, but also ignoring potentially relevant possible scenarios. For example, lineages that have entered India from Madagascar before their breakup, could have been evolving in relative isolation there until coming into contact with Eurasia. However, we suspect that testing that kind of scenario would require more detailed geographical resolution, and thus may warrant a dedicated separate study.

Each taxon’s range was defined as the one or several areas it occurred in, based on the collected information from GBIF and the literature. Where information on continent was missing, it was inferred from coordinates and countries of occurrence, and inconsistencies between these were checked and corrected. Taxa occurring in many continents were inspected for the extent of overlap and reliability of the records. In doing so, we also paid attention to species that were introduced to areas by humans, and removed such occurrences from the species’ range, thereby only assigning its presumed natural range.

Given the potential age of the group, it is conceivable that tectonic plate movement played a role in their dispersal and distribution. We thus constructed a stratified dispersal matrix representing the changing strength of dispersal barriers between the continents over time. We divided the last 200ma before the present into five time slices, following the tectonic events described in Seton *et al*. (2012), stretching from 200-150ma (Pangaea intact), 150-110ma (breakup of Pangaea into Gondwana and Laurasia, Madagascar breaking off of Africa, though still connected via Antarctica), 110-50ma (breakup of Gondwana into Africa, Australia-Antarctica, and South America, the latter still connected to Antarctica via a land bridge), 50-20ma (Australia separates from Antarctica, Laurasia breakup into Laurentia and Eurasia (Hosner, Braun & Kimball 2015), South America properly disconnected from Antarctica (Reguero *et al*. 2014)), and 20-0ma (land bridges (some temporary) establish at Beringia (Hosner, Braun & Kimball 2015), the Isthmus of Panama, and between Australia and Asia, and Eurasia and Africa).

Inspired by Toussaint, Bloom and Short (2017), we recognized dispersal barriers of different strengths: 1) directly adjacent areas (barrier of strength 0), 2) areas connected by land bridge (0.15), 3) areas separated by a small distance of water (0.25), and 4) by a large distance of water (0.75). Having to pass through another area was considered a barrier of strength 0.5, and the case of facing more than 3 barriers was assigned a strength of 0.95; the corresponding dispersal multiplier was 1 - strength of barriers. The route of smallest resistance was picked for each possible dispersal case, adding up the different barriers encountered (matrices see Table S4).

While it would make sense that different dispersal barriers at different times would affect the dispersal dynamics of the group, it is also possible that only time differences or only dispersal barriers did. Furthermore, the partially arbitrary choice of time intervals could affect the inference as well. Thus, we also calculated averaged dispersal matrices to be used without time stratification. For this purpose, the dispersal multipliers of each time slice were weighted by the duration of that time slice relative to the total time from the beginning of that slice until the present. These weighted multipliers were added up for each transition and then divided by the sum of weights, so they would add to one. This weighting is intended to represent the fact that the more recent positioning of the continental plates should have had a higher impact on the current distribution of extant taxa. Two matrices (for 3 and 4 time slices respectively, see Table S5) were constructed that way, to be used for either the phylogeny dated with the younger or older calibration times.

We used the package *BioGeoBEARS* (Matzke 2013b; Matzke 2013a) to estimate the ancestral ranges of the dung beetles. The package implements some of the most popular models for ancestral range estimation, DEC (Ree & Smith 2008), DIVA (Ronquist 1997), and BayArea (Landis *et al*. 2013), in the same framework, allowing them to easily be compared against each other to test different biogeographical hypotheses. While the DEC model is a maximum likelihood implementation just as originally implemented in Lagrange, the DIVA model was originally implemented using parsimony, and in *BioGeoBEARS* the processes DIVA assumes are modeled under maximum likelihood. Similarly, BayArea was originally implemented as Bayesian, and is represented in *BioGeoBEARS* as a maximum likelihood interpretation of the same. Thus, these two models should be referred to as DIVALIKE and BAYAREALIKE respectively.

A popular feature of *BioGeoBEARS* is the addition of “jump-dispersal” to any of these models. By adding the additional jump parameter (“+j”), one allows for founder events in the model, meaning that at a speciation event, one descending lineage stays in the ancestral range, while the other descendant jumps to a new area (Matzke 2014). However, there has recently been some debate regarding the validity of the +j models (Ree & Sanmartín 2018; Klaus & Matzke 2019), and this type of cladogenetic event is believed to be more important in island systems than in non-island clades (Matzke 2013b). For those reasons, we decided not to employ the j parameter.

However, since the dispersal multiplier matrices defined above are somewhat arbitrary, we do pair them with a parameter (w) that scales the matrix and that can be optimized as well, constituting the +w models (Dupin *et al*. 2017). The parameter w is used to exponentiate the dispersal multiplier before it is multiplied by the dispersal rate and is 1 by default. Thus, if the multipliers play an important role in the biogeographical history of the group, w is likely to be estimated to be larger than 1, whereas it should be estimated to be lower than 1 if the dispersal matrix does not add much to explain the patterns. While this does not allow to modulate the relative strengths of the different multipliers, it seems reasonable to expect that w would be estimated to downweigh the importance of a grossly unrealistic set of multipliers.

We employed each of the three base models (DEC, DIVALIKE, BAYAREALIKE) in four different ways: 1) as a basic model (estimating d and e), 2) with the non-stratified manual dispersal multiplier matrix and estimated w parameter, 3) as a time stratified model, and 4) as a time stratified model with a manual dispersal multiplier matrix for each time slice, and estimated w parameter. In all those analyses, we constrained the maximal range any lineage can occupy to 3, as none of the extant species occupy more than 3 areas. To account for the branch support issues, we ran all these models both on the trees with old and young calibrations, as well as the ten bootstrap trees each.

Finally, preliminary analyses produced curious results, particularly in the time stratified model, where lineages within clades that where entirely present in one area (*e*.*g*. Madagascar) would commonly disperse to a neighboring area (*e*.*g*. Africa) right after a speciation event, only to return to the ancestral area again. It was presumed that clades in which every species inhabited the same two areas (*e*.*g*. the Americas) would be responsible for this, as models as DEC do not include the required scenario (cladogenesis in widespread taxa where both daughters inherit widespread range). Thus, such cases required the above pattern of one daughter losing and re-gaining the widespread range, thereby inflating the inferred dispersal rate and forcing other lineages to do the same. We therefore created test data sets where we identified the area in which species most commonly co-occurred (the Americas), or the two pairs of most co-inhabited areas (the Americas and Eurasia), and coded those as one, thus getting a 6-area and 5-area dataset to be analyzed separately.

### Diversification Analysis

We tested two sets of biogeography-related hypotheses: whether areas with larger mammals and more diverse droppings are associated with higher diversification rates of dung beetles, and whether diversification rates were raised when dung beetles gained access to new habitats, dispersing from Gondwana to Madagascar or Laurasia. The former hypothesis was derived from Davis and Scholtz (2001); (also elaborated in Scholtz, Davis & Kryger 2009). They classify mammalian dung into four types based on size and physico-chemical characteristics: 1) small dry pellets from small to medium herbivores, 2) small odiferous droppings from omni- or carnivores, 3) large, dry, course-fibered droppings from large non-ruminant herbivores, and 4) large, moist, fine-fibered pads from large ruminants. The number of those types of dung available, as well as the (fairly correlated) body size of mammals was tallied for different biogeographical regions, and shown to relate to aspects of dung beetle diversity (Davis & Scholtz 2001; Scholtz, Davis & Kryger 2009).

Explicitly biogeographic diversification models such as GeoSSE (Goldberg, Lancaster & Ree 2011) or GeoHiSSE (Beaulieu & O’Meara 2016; Caetano, O’Meara & Beaulieu 2018) only test diversification between two areas (and three ranges: endemic to one or the other area, or widespread), making explicitly testing hypotheses involving more areas (not to speak of all seven) impossible. However, in the case of only three areas, testing different combinations of two against each other can still yield relevant insights. To this end, we re-coded the distribution data from above to merge occurrences 1) of areas with large mammals and diverse droppings (Afro-Eurasia: Africa, Europe, and Asia), of medium sized mammals and less diverse droppings (North and South America), and those of those with small mammals and the least diverse droppings (East Gondwanan Fragments: Oceania and Madagascar).; and 2) those of the areas formerly making up Gondwana (South America, Africa, and Oceania), those of former Laurasia (North America, Europe, and Asia), while keeping Madagascar as an area of its own. We then formatted these two data sets to suit the different GeoSSE tests: a set where we join the areas with medium sized mammals and intermediate droppings-diversity with the areas of large mammals and diverse droppings, and one where we join them with the small mammal and low droppings-diversity instead. Another set of where we join Madagascar with Gondwana, where we join it with Laurasia, and where we leave it as an area separate from the rest.

For each of the resulting five data sets (and for both the young and old tree respectively), we inferred maximum likelihood estimates with GeoSSE, under a set of different model constraints. The constraints used were combinations of equal speciation between areas, equal extinction between areas, equal dispersal rates between areas, and variation where speciation was set to zero in one area and forced to be equal in the other and widespread lineages, and vice versa. The full and constrained models were then compared using likelihood ratio tests and their ΔAIC score. The best models for each combination of data set and tree were subsequently used to get posterior distributions of the parameter estimates using MCMC, with an exponential prior related to the Kendall-Moran estimate for net diversification rate (Kendall 1949; Moran 1951). A short preliminary chain of 100 generations was run and the distances between the 5% and 95% quantiles for each parameter were used to set the tuning parameter w for the slice sampler. Then, each dataset was run for 20,000 generations. Convergence was assessed by the convergence parameter of the function, by visual inspection of the log likelihood trace, and calculating effective sample size using *effectiveSize* from the R package *coda* (Plummer *et al*. 2004). We then compared the 95% quantiles of the posterior distribution for all estimated parameters to see whether they overlap.

## Results

### Phylogeny and Divergence Times

Out of 122 genera represented in this phylogeny, 33 were monophyletic, 28 were non-monophyletic, and 61 were monotypic (Table S2). When considering the bootstrap trees, 14 genera were consistently monophyletic, 22 consistently non-monophyletic, whereas the remaining 25 varied between trees. On the tribal level, we recovered Eucraniini, Ateuchini, Eurysterniini, Gymnopleurini, and Sisyphini as monophyletic (Table S3). Of these, only the last three consistently so, with Onitini being monophyletic in some bootstrap replicates. Upon closer inspection of the reasons for each tribe’s non-monophyly (Table 1), we see that at least part of it results from taxonomic issues. For Scarabaeini, Phanaeini, and Onitini, non-monophyly is due to a few *incertae sedis* taxa, most of which monotypic. Oniticellini and Onthophagini have similar issues where one intruder of the former is a member of the latter, but also the whole of Oniticellini is nested within Onthophagini. Dichotomiini, Deltochilini and Canthonini are scattered in clumps across the tree, with the latter two intermingling a lot, while Coprini come out in three separate clades.

The estimated ages of the calibrated nodes for both the young and old calibration are given in Table 2. It is apparent that under both calibration schemes, the crown age of the Scarabaeinae has hit its constrained maximum age, while the stem ages of the constrained nodes are not reaching their maximum constraints but are often meeting or exceeding the minimum age constraint of the subfamily’s crown. The bootstrap branch support values can be seen in Figure 1.

**Figure 1:**
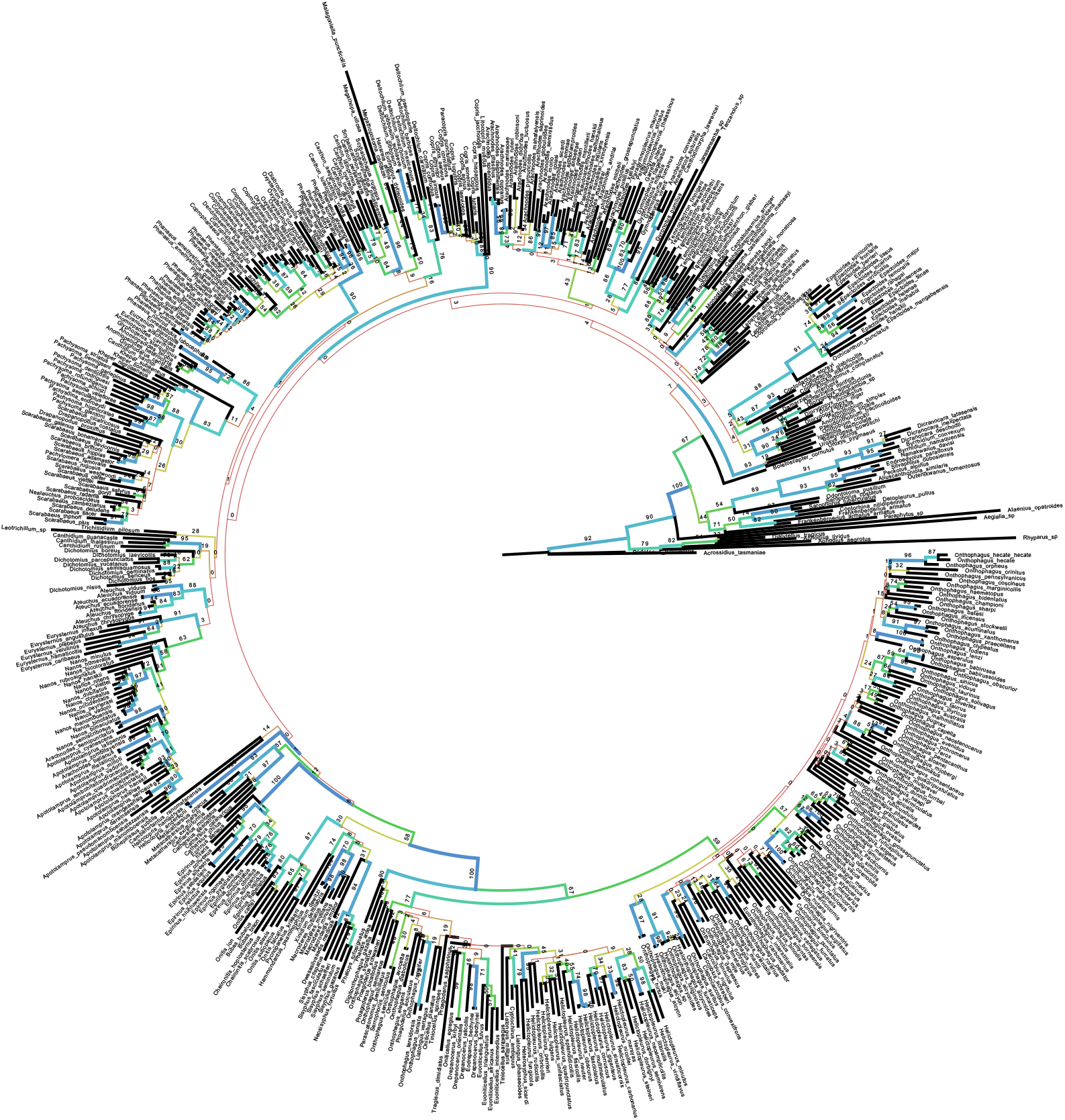
Branch Support Dung Beetle Phylogeny. Bootstrap branch length support on the un-dated RAxML phylogeny. Colder colors indicating higher branch support, decreasing branch width with decreasing support (terminal branches are black).

### Ancestral Range Estimation

For all analyses of the 7-area data set on the young and old maximum likelihood trees and their respective sets of 10 bootstrap trees, the analyses recovered the unstratified DEC model with manual dispersal multipliers (DEC+w) as the best-fitting model. In all trees, w – the exponent for the dispersal multipliers – was estimated to be larger than 1 (1.71-2.83), indicating its weight in the dispersal process being higher than the initial dispersal matrix suggested. The estimated dispersal and extirpation rates vary between the trees but are comparable and at the very least in the same order of magnitude. For the 6-area and 5-area datasets, which model was inferred to fit best varied widely among the different trees.

The ancestral range of the whole subfamily was estimated to be Africa, Oceania, and South America for both the young and old tree. However, in the alternative trees based on bootstrap replicates, the estimated ancestral range could include Madagascar instead of South America or instead of Oceania, or could just include Africa and Oceania (Table S6). Interestingly, reconstructions with root states other than Africa, Oceania, and South America tended to be more ambiguous. Overall, the estimated ancestral ranges seemed to suggest that numerous clades mostly stay in the same areas (particularly Africa, Oceania, and Madagascar), with some more dispersal within the Americas and Eurasia, as well as within the clade comprising the Onthophagini and Oniticellini (Figure S1).

### Diversification Analyses

The best fitting models for each data set and tree, according to likelihood ratio tests and ΔAIC scores, are given in Table 3. The MCMC analyses converged and yielded reasonably high effective sampling sizes with no value below 295. The 95% quantiles of the posterior distributions for each parameter estimate is given in Table 4.

**Table 3.**
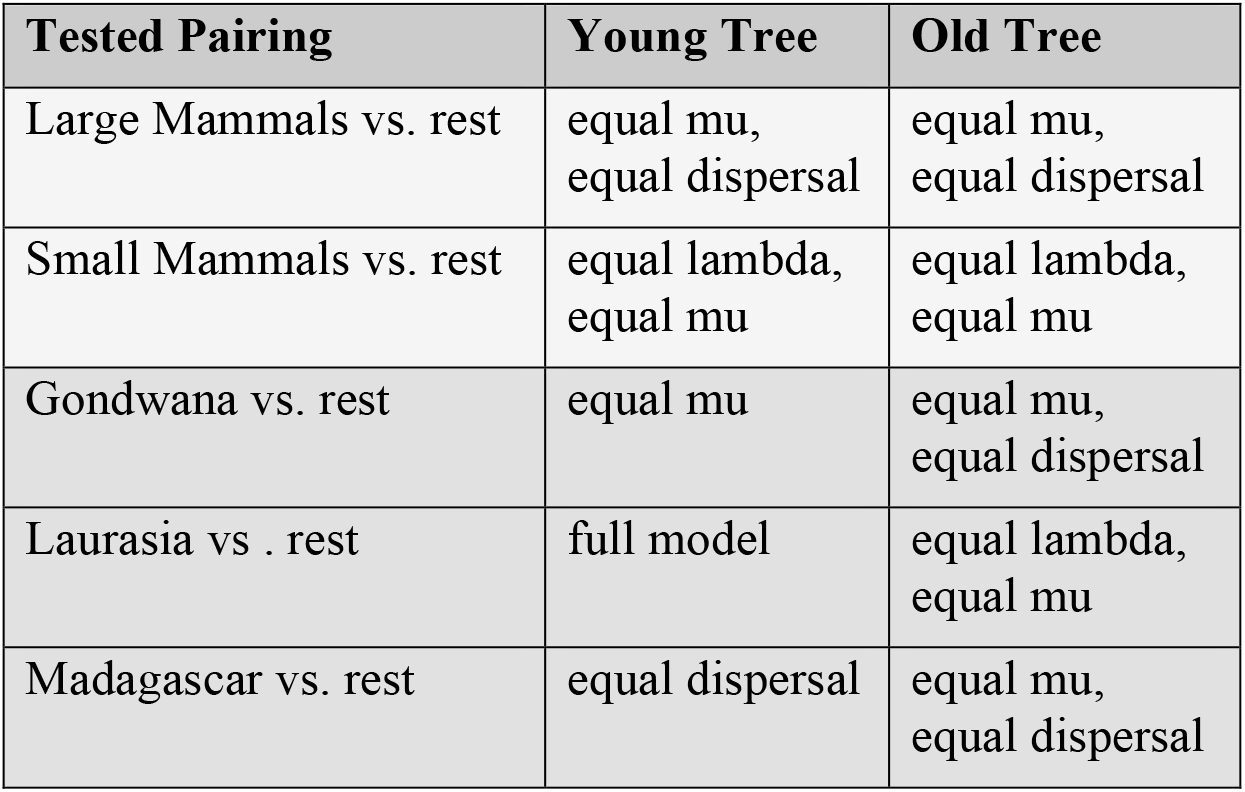
Best Fitting Models for Biogeographcal Diversification Hypotheses. Best fitting models of ML GeoSSE analyses for each data set and each tree, as determined by likelihood ratio tests and ΔAIC scores. Models where lambda was constrained to be equal in both areas, also constrained lambda to be zero for widespread taxa.

| Tested Pairing | Young Tree | Old Tree |
| --- | --- | --- |
| Large Mammals vs. rest | equal mu,<br>equal dispersal | equal mu,<br>equal dispersal |
| Small Mammals vs. rest | equal lambda,<br>equal mu | equal lambda,<br>equal mu |
| Gondwana vs. rest | equal mu | equal mu,<br>equal dispersal |
| Laurasia vs . rest | full model | equal lambda,<br>equal mu |
| Madagascar vs. rest | equal dispersal | equal mu,<br>equal dispersal |

**Table 4.** Posterior Distribution of GeoSSE Rate Estimates. 95% quantiles of posterior distributions for diversification and dispersal rates estimated under the best scoring model for each data set and each tree. s=speciation rate, x=extinction rate, r=net diversification rate (s-x), d= dispersal rate.

| Pairings | Quant. | sA | sB | sAB | xA | xB | rA | rB | rAB | dA | dB | Likelihood |
| --- | --- | --- | --- | --- | --- | --- | --- | --- | --- | --- | --- | --- |
| Young Tree |  |  |  |  |  |  |  |  |  |  |  |  |
| Only Large Mam. Vs. rest | 2.50% | 0.0968 | 0.0942 | 0.0382 |  | 0.0192 | 0.0636 | 0.0600 | 0.0382 |  | 0.0008 | -2436.8114 |
|  | 97.50% | 0.1449 | 0.1437 | 3.9306 |  | 0.0785 | 0.0803 | 0.0799 | 3.9306 |  | 0.0021 | -2428.7662 |
| Only Small Mam. vs. rest | 2.50% |  | 0.1076 |  |  | 0.0342 |  | 0.0613 |  | 0.0002 | 0.0015 | -2436.8855 |
|  | 97.50% |  | 0.1552 |  |  | 0.0922 |  | 0.0756 |  | 0.0014 | 0.0042 | -2431.4368 |
| Madagascar vs. rest | 2.50% | 0.1041 | 0.0930 | 0.1085 | 0.0006 | 0.0191 | 0.0770 | 0.0602 | 0.1085 |  | 0.0003 | -2381.0849 |
|  | 97.50% | 0.2102 | 0.1423 | 8.0402 | 0.1276 | 0.0798 | 0.1158 | 0.0756 | 8.0402 |  | 0.0010 | -2373.0650 |
| Laurasia vs . rest | 2.50% | 0.0819 | 0.0863 | 0.0016 | 0.0006 | 0.0160 | 0.0581 | 0.0528 | 0.0016 | 0.0223 | 0.0003 | -2619.1510 |
|  | 97.50% | 0.1308 | 0.1409 | 0.0651 | 0.0674 | 0.0857 | 0.0928 | 0.0724 | 0.0651 | 0.0577 | 0.0036 | -2610.9780 |
| Gondwana vs. rest | 2.50% | 0.0964 | 0.0683 | 0.0002 |  | 0.0008 | 0.0863 | 0.0593 | 0.0002 | 0.0051 | 0.0028 | -2629.9069 |
|  | 97.50% | 0.1228 | 0.0924 | 0.0309 |  | 0.0305 | 0.1054 | 0.0721 | 0.0309 | 0.0156 | 0.0056 | -2622.8756 |
| Old Tree |  |  |  |  |  |  |  |  |  |  |  |  |
| Only Large Mam. vs. rest | 2.50% | 0.0636 | 0.0619 | 0.0286 |  | 0.0093 | 0.0448 | 0.0426 | 0.0286 |  | 0.0005 | -2632.4797 |
|  | 97.50% | 0.0952 | 0.0942 | 1.5142 |  | 0.0486 | 0.0560 | 0.0557 | 1.5142 |  | 0.0013 | -2624.6484 |
| Only Small Mam. vs. rest | 2.50% |  | 0.0698 |  |  | 0.0183 |  | 0.0435 |  | 0.0001 | 0.0010 | -2633.0250 |
|  | 97.50% |  | 0.0994 |  |  | 0.0546 |  | 0.0530 |  | 0.0009 | 0.0029 | -2627.6812 |
| Madagascar vs. rest | 2.50% | 0.0726 | 0.0575 | 0.0952 |  | 0.0043 | 0.0558 | 0.0437 | 0.0952 |  | 0.0002 | -2578.7214 |
|  | 97.50% | 0.1066 | 0.0887 | 6.9666 |  | 0.0438 | 0.0753 | 0.0544 | 6.9666 |  | 0.0007 | -2571.5387 |
| Laurasia vs . rest | 2.50% |  | 0.0602 |  |  | 0.0103 |  | 0.0394 |  | 0.0137 | 0.0002 | -2810.4804 |
|  | 97.50% |  | 0.0851 |  |  | 0.0443 |  | 0.0513 |  | 0.0327 | 0.0027 | -2805.3483 |
| Gondwana vs. rest | 2.50% | 0.0627 | 0.0477 | 0.0001 |  | 0.0003 | 0.0580 | 0.0437 | 0.0001 |  | 0.0028 | -2829.6036 |
|  | 97.50% | 0.0780 | 0.0611 | 0.0174 |  | 0.0152 | 0.0702 | 0.0512 | 0.0174 |  | 0.0045 | -2822.8190 |

Regardless of whether the areas with medium sized mammals and intermediate diversity in droppings were joined with the areas of large or small mammals, there was no significant difference between them in terms of diversification rates (if the best model did not constrain them; Figure 2, Figure 3). When the areas of medium and small mammals are joined, the analysis attributes any difference in diversity between them to higher dispersal out of Afro-Eurasia into the other areas.

**Figure 2:**
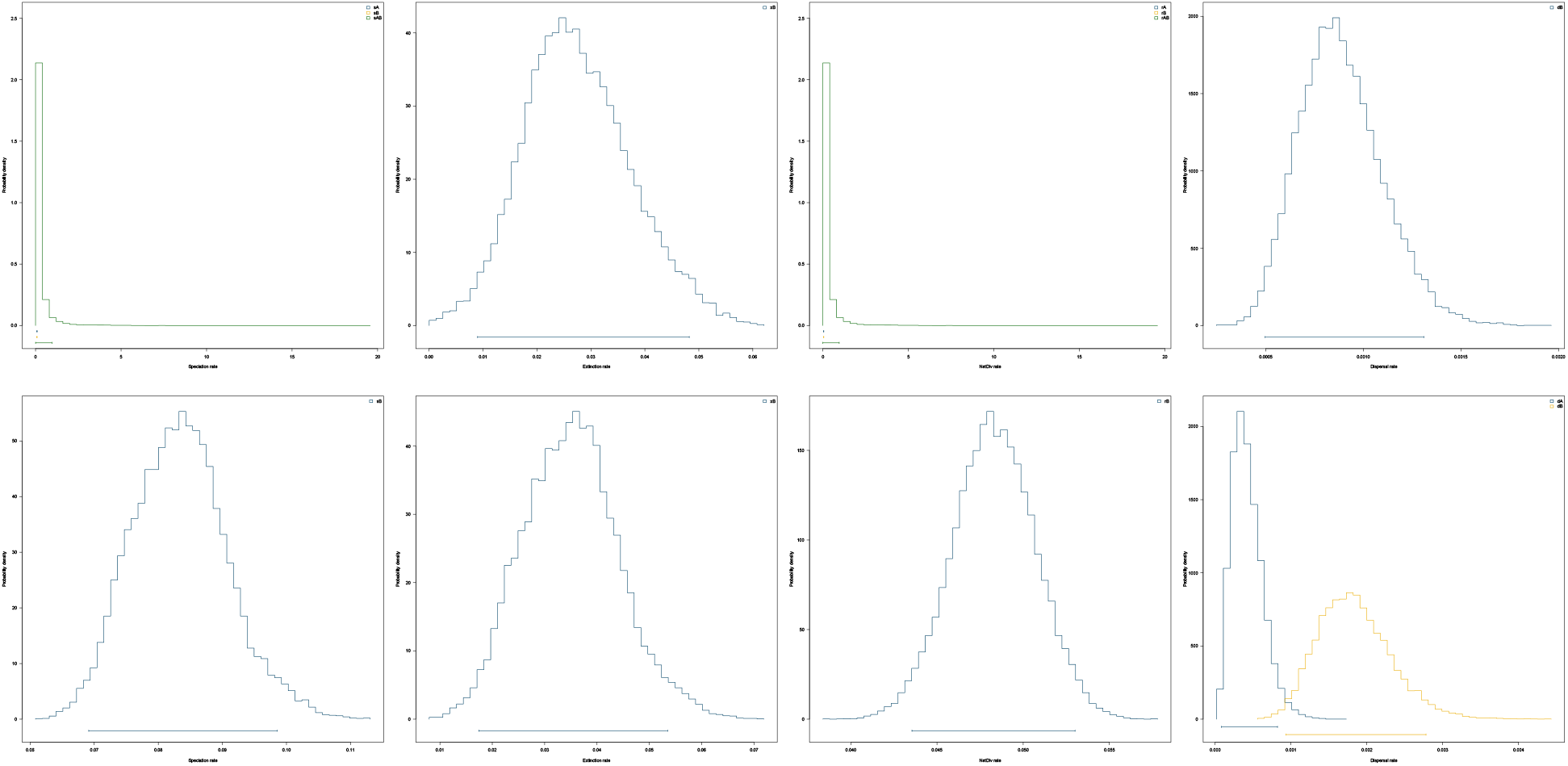
GeoSSE Results Dung Producers Old Tree. Posterior distributions of area-dependent rates estimated by GeoSSE. Left to right: speciation rates, extinction rates, net diversification rates (speciation – extinction), dispersal rates. **Top**: rates in areas with large and medium sized mammals and high and intermediate droppings-diversity (Afro-Eurasia and Americas, blue), and rates in areas with small sized mammals and low droppings-diversity (East Gondwanan Fragments, yellow). **Bottom**: rates in areas with large sized mammals and high droppings-diversity (Afro-Eurasia, yellow), and rates of areas with low and medium sized mammals and low and intermediate droppings-diversity (Americas and East Gondwanan Fragments, blue). Rates of widespread taxa are colored green; where only one rate was estimated, it is colored blue. Dispersal rates of an area reflect dispersal *out* of said area.

**Figure 3:**
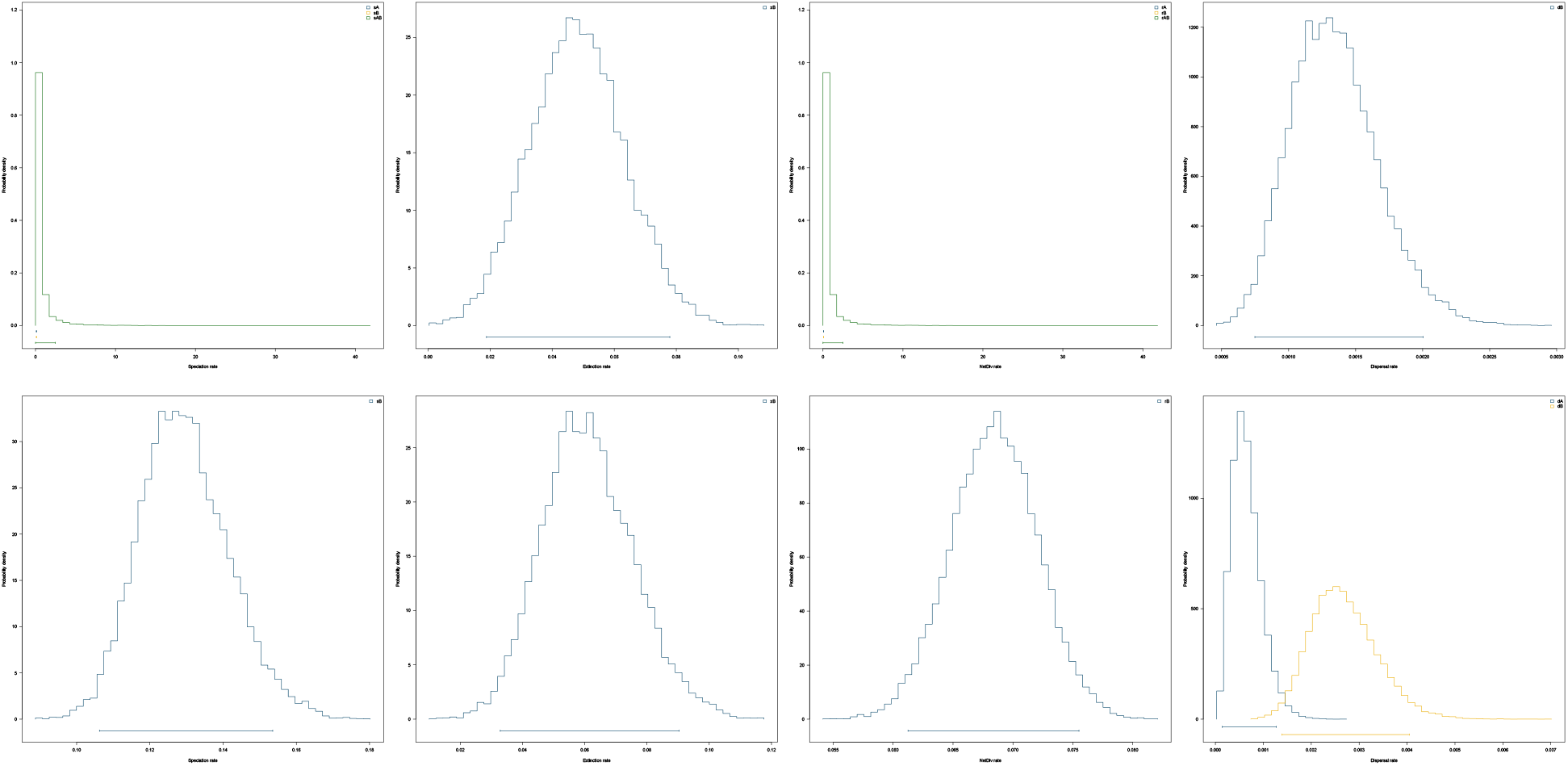
GeoSSE Result Dung Producers Young Tree. Posterior distributions of area-dependent rates estimated by GeoSSE. Left to right: speciation rates, extinction rates, net diversification rates (speciation – extinction), dispersal rates. **Top**: rates in areas with large and medium sized mammals and high and intermediate droppings-diversity (Afro-Eurasia and Americas, blue), and rates in areas with small sized mammals and low droppings-diversity (East Gondwanan Fragments, yellow). **Bottom**: rates in areas with large sized mammals and high droppings-diversity (Afro-Eurasia, yellow), and rates of areas with low and medium sized mammals and low and intermediate droppings-diversity (Americas and East Gondwanan Fragments, blue). Rates of widespread taxa are colored green; where only one rate was estimated, it is colored blue. Dispersal rates of an area reflect dispersal *out* of said area.

Despite some differences in which model fitted the Gondwanan origin data best, the results between the old tree (Figure 4) and the young tree (Figure 5) are largely consistent. They show that lineages in Gondwana have a significantly lower speciation rate than lineages outside of it, while lineages in Laurasia disperse into the other areas at a higher rate than vice-versa, while there are no significant differences in either diversification nor dispersal between Madagascar and the other areas.

**Figure 4:**
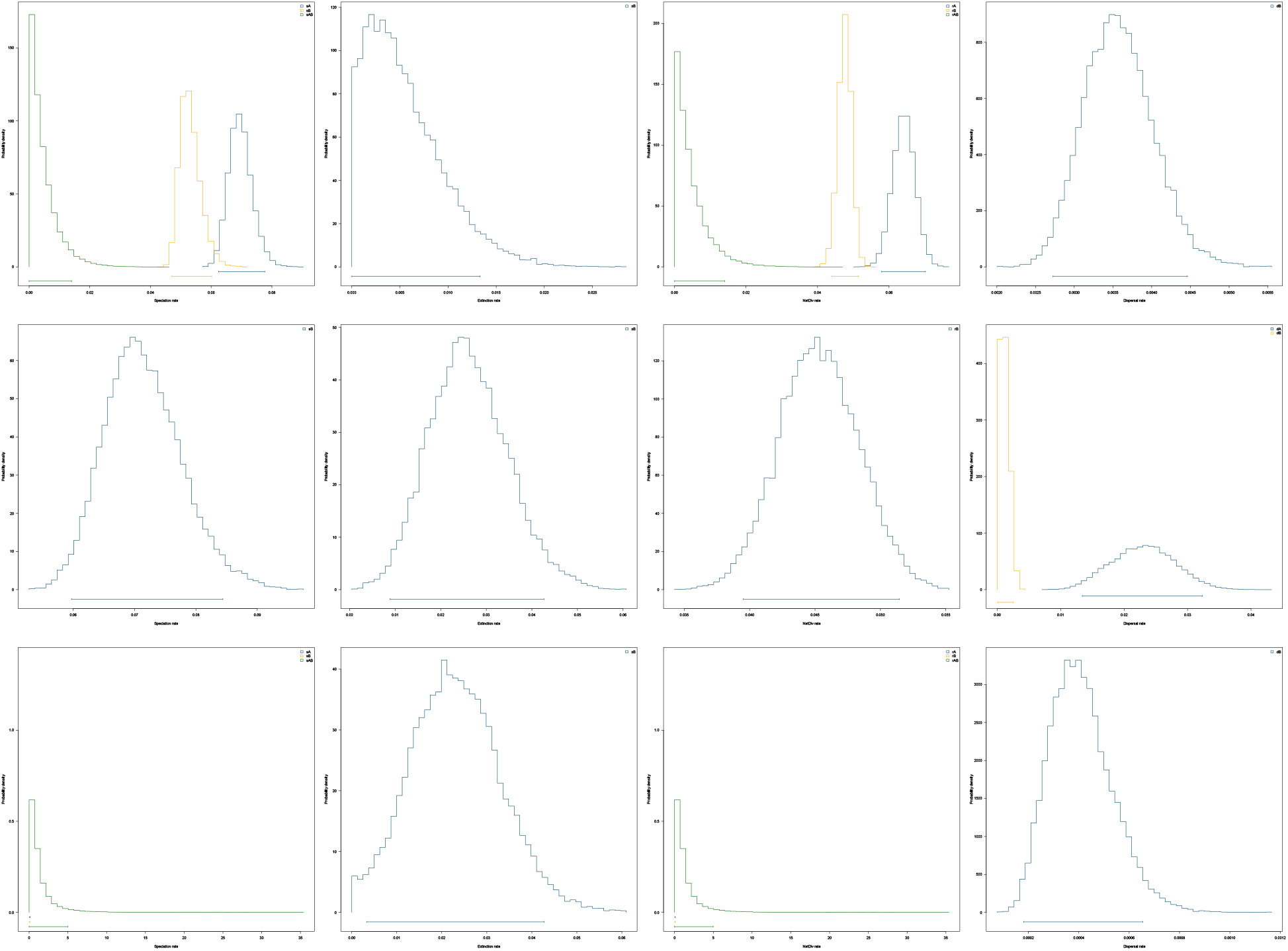
GeoSSE Results Out-Of-Gondwana Old Tree. Posterior distributions of area-dependent rates estimated by GeoSSE. Left to right: speciation rates, extinction rates, net diversification rates (speciation – extinction), dispersal rates. **Top**: rates in Laurasia and Madagascar (blue), and rates in Gondwana (yellow). **Center**: rates in Laurasia (blue), and rates in Gondwana and Madagascar (yellow). **Bottom**: rates in Madagascar (blue), and rates in Gondwana and Laurasia. Rates of widespread taxa are colored green; where only one rate was estimated, it is colored blue. Dispersal rates of an area reflect dispersal *out* of said area.

**Figure 5:**
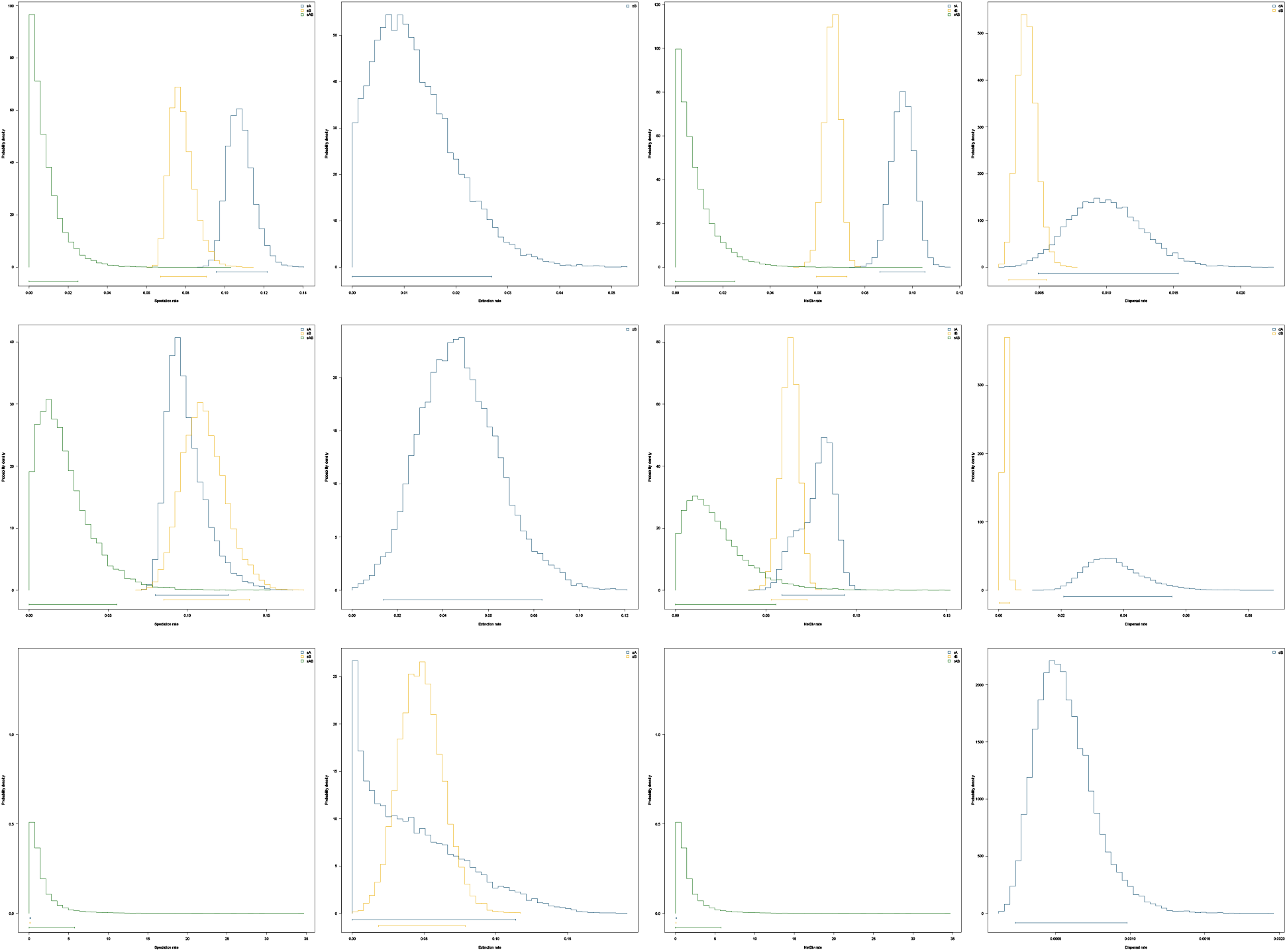
GeoSSE Results Out-Of-Gondwana Young Tree. Posterior distributions of area-dependent rates estimated by GeoSSE. Left to right: speciation rates, extinction rates, net diversification rates (speciation – extinction), dispersal rates. **Top**: rates in Laurasia and Madagascar (blue), and rates in Gondwana (yellow). **Center**: rates in Laurasia (blue), and rates in Gondwana and Madagascar (yellow). **Bottom**: rates in Madagascar (blue), and rates in Gondwana and Laurasia. Rates of widespread taxa are colored green; where only one rate was estimated, it is colored blue. Dispersal rates of an area reflect dispersal *out* of said area.

## Discussion

### Phylogeny and Taxonomy

With 541 represented in-group species, this is to date the most inclusive dated species-level molecular phylogeny of Scarabaeinae. Most previously inferred phylogenies of the group were either constrained to specific sub-clades or regions (e.g. Davis & Scholtz 2001; Wirta, Orsini & Hanski 2008; Sole & Scholtz 2010; Wirta *et al*. 2010; Mlambo, Sole & Scholtz 2015; Breeschoten *et al*. 2016; Gunter *et al*. 2018), or part of a larger phylogeny where the main focus accordingly was not or not only on dung beetles (Ahrens, Schwarzer & Vogler 2014; Kim & Farrell 2015; Gunter *et al*. 2016; Toussaint *et al*. 2017). A recent large un-dated phylogeny by Tarasov and Dimitrov (2016) based on 8 gene regions had a similar amount of terminals (547 with outgroup), though it constitutes a smaller sample of actual species diversity, as many species were represented by multiple accessions and many were not determined to species level. The levels of generic and tribal monophyly in this new phylogeny (Table S2, Table S3) and the causes of it (Table 1) would suggest a reasonable level of agreement between this tree and current taxonomy. The lack of support for many groupings within the tree (Figure 1), particularly in the backbone, is cause for concern, as it not only suggests shortcomings in the inferred tree, but also casts doubt upon the reliability of analyses results derived from that tree. Other attempts at phylogenies of the overall Scarabaeinae were plagued with similar patterns of low branch support (Tarasov & Dimitrov 2016), which suggests a general issue in the study of this group. Molecular phylogenies have challenged the traditional classification, particularly the tribal monophyly of Canthonini and Dichotomiini (Monaghan *et al*. 2007), as well as Coprini, Onthophagini and Oniticellini, while the monophyly of the remaining tribes still seems supported (Scholtz, Davis & Kryger 2009). Tarasov and Dimitrov (2016) note how their own results are consistent with those of previous phylogenetic studies of Scarabaeinae (Ocampo & Hawks 2006; Monaghan *et al*. 2007; Vaz-de-Mello 2007; Wirta, Orsini & Hanski 2008; Sole & Scholtz 2010; Wirta *et al*. 2010; Mlambo, Sole & Scholtz 2014; Gunter *et al*. 2016), as well as with a large phylogeny based on morphology (Tarasov & Génier 2015). They furthermore observed that the studies to date tend to resolve old nodes and more recent nodes, but not intermediate ones, and that the same set of problematic tribes mentioned above are consistently not monophyletic. All of this seems to be reflected in our results as well, particularly when considering the extent of monophyly problems (Table 1). Using the suggested new classification by Tarasov and Dimitrov (2016) does not yield much improvement. While Eucraniini and Eurysternini are strictly monophyletic in both, monophyly gained through reclassification in Dichotomiini, Canthonini, and Scarabaeini is offset by the loss of it in Ateuchini, Gymnopleurini, and Sisyphini. Both classifications show similar consistency across the set of bootstrap trees.

### Ancestral Range Estimation and the Origin of Scarabaeinae

The two main hypotheses regarding where dung beetles originated are an origin in Gondwana followed by vicariance events after the breakup of the supercontinent (Hanski & Cambefort 1991; Davis & Scholtz 2001; Davis, Scholtz & Philips 2002), and an origin in Africa and subsequent dispersal out of it (Sole & Scholtz 2010). One major point of conflict between those two ideas was the age of Scarabaeinae: Gondwanan vicariance would necessitate the group to be of Mesozoic, rather than Cenozoic origin, as they would have to exist and be widespread enough before the continental breakup (110ma according to Sanmartin and Ronquist (2004), 93ma according to Scotese (1993)) in order for vicariance events to be plausible.

To answer the question of biogeographic origin in absence of an appropriate phylogeny, some workers relied on classification (Hanski & Cambefort 1991; Davis & Scholtz 2001), considering widespread tribes to be ancient and predating the Gondwana-breakup, in turn giving rise to younger, less widespread tribes. Later attempts combined the relative age of phylogenies (Monaghan *et al*. 2007; Wirta, Orsini & Hanski 2008) with fast and slow rates of molecular sequence divergence in insects to get maximal and minimal divergence time estimates (Scholtz, Davis & Kryger 2009), concluding that even the slowest known divergence rates would not support the idea of a pre-Gondwanan-breakup origin of dung beetles. Sole and Scholtz (2010) subsequently used a time calibrated phylogeny of the African representatives of Canthonini and Dicotomiini to address the question, finding the divergence times between dung beetles and their outgroup, and of the crown of dung beetles to be considerably younger than the breakup of Gondwana (56ma and 40ma respectively), thus further supporting the later out-of-Africa scenario.

As for the divergence times inferred in this study (Table 2), the older age of 131.6ma would place the origin of dung beetles well before the complete breakup of Gondwana, whereas the younger estimate of 98.1ma appears ambiguous, depending on when the actual separation of Gondwana was completed, and depending on the accuracy of this age estimate. In either case however, the inference of older origins, similar to other recent estimates that considered more than just a subset of Scarabaeinae (Ahrens, Schwarzer & Vogler 2014; Gunter *et al*. 2016), makes a Gondwanan origin and thus the potential for vicariance after its breakup seem like a plausible option again.

The reconstruction of what is essentially Gondwana as the ancestral range at the crown of our phylogenies seems to further support the idea of Gondwanan-vicariance. However, the DEC model is known to have a bias towards widespread ancestors (Clark *et al*. 2008; Ree & Smith 2008; Buerki *et al*. 2011; Matzke 2014), even if not as strongly as DIVA (Kodandaramaiah 2010). Therefore, since we constrained the number of areas a lineage can maximally inhabit to three, the inference of those three areas as the origin could possibly be an artefact. With regards to the branch support issues, it would appear that the consistency with which the same model was preferred across all trees using the 7-area dataset, and the relative stability of the inferred origin to be Gondwana, or parts thereof (Table S6), could be seen as a sign that the result is robust enough. However, the wildly varying best supported model under the reduced area datasets (6 and 5 areas) is cause for concern. It could be argued, that leaving those pairs or areas separate (North and South America, Europe and Asia respectively), might lead to a distribution of areas across clades that is not broken up when they are rearranged in the bootstrap trees, thus implying the same dispersal mechanisms. On the contrary, lumping them could lead to single-distribution clades being broken up, thus changing the number of implied dispersal and vicariance events. However, this is rather speculative and requires further investigation. Finally, while the high estimates for w suggest that the specified dispersal multipliers are a relevant improvement of the model overall, this does not guarantee that they are an accurate representation of dung beetle dispersal probabilities at the given times. While they might capture some large-scale dispersal constraints, the relative magnitude of the dispersal multipliers between specific continents could still be inaccurate, *e*.*g*. because of the way the barriers were specified and the values they were assigned. Sensitivity tests, or even adding an approach using actual distance, could help to inform us about this.

In any case, given the branch support issues, we would refrain from a too detailed interpretation of the ranges reconstructed at internal nodes. However, it is notable that the reconstruction suggests that many clades can be found which seem to be predominantly confined to one (or few) areas, particularly to Oceania, Madagascar, and Africa, with some more dispersal in American or Eurasian clades. An exception seems to be the clade comprising Onthophagini and Oniticellini, which seems to have much more area changes. This could indicate that the Onthophagini and Oniticellini are different from the remaining dung beetles in their biogeographic history and maybe their dispersal abilities. It could however also just reflect a lack of sampling in that clade.

### Diversification Analyses

The GeoSSE results for the hypothesis on whether mammal size and available dung-diversity in different biogeographical areas affected the diversification rates of Scarabaeinae seem to suggest that this is not the case. The increased dispersal from Afro-Eurasia into the other regions is certainly interesting and could reflect different plausible dispersal events. But given that the test was not set up to address these alternatives, we would suggest exercising care not to overinterpret this pattern. When Davis and Scholtz (2001) reported on the patterns among mammals, the diversity of their droppings, and dung beetle diversity in different areas of the world, they noted that it was particularly related to tribal diversity (and generic diversity within tribes) in those areas, but less correlated to genus or species diversity. They interpret this as a sign that the influence of these patterns in mammal body size and droppings on dung beetle diversification happened earlier in their evolutionary history, around the time when the tribes split. While the idea that mammal dung availability could have been a crucial influence on beetle diversification in the past appears very sensible, it would seem that the arguments presented in this particular case are problematic. Firstly, it relates an extant pattern in mammals to past effects in beetles assuming that the distribution of mammals and their droppings across the world was comparable between then and now. However, those extant patterns were almost certainly not constant over the timespan of dung beetle diversification, with the most recent event that changed those patterns already being the Late Quaternary megafaunal extinctions (Stuart 2015). Furthermore, assuming from the ages in this current tree (Table 2), as well as past estimates (e.g. Gunter *et al*. 2016), the ages of many tribes may either be older than the rise of mammal diversity, or would at least coincide with a time when mammal diversity would not be expected to compare well to the extant one. Finally, considering the fact that the pattern does not correlate well with species level diversity, the method we employed might also be suboptimal to test the implied scenario of past influence, and an approach that allows the influence of mammal size and droppings to be constrained to particular time-intervals might be more suitable.

The result of the Gondwanan origin GeoSSE analysis might initially seem surprising, as a large portion of Scarabaeinae species diversity is found in areas that were formerly Gondwana (Scholtz, Davis & Kryger 2009) and the former-Gondwanan lineages were not underrepresented in comparison to the others (Gondwana: 320 taxa, Laurasia: 143 taxa, Madagascar: 114 taxa). But considering the possible Gondwanan origin of the whole group, and the perceived inertia of clades (diversifying in an area rather than dispersing more), one might suggest that the larger absolute number of species in former Gondwanaland results from diversifying at a lower rate but over longer timespans than the on average probably younger Laurasian or Malagasy lineages. This would also explain the lack of rate difference between the latter and Laurasia and Gondwana combined. The higher dispersal out of Laurasia was not expected, assuming the general direction of dung beetle dispersal was outward from Gondwana. But in the light of higher dispersal outside of Gondwana, this could reflect the dispersal of few lineages out of Gondwana, where they diversified, and subsequently a few of those Laurasian taxa returned to former Gondwana (*e*.*g*. in the case of taxa which returned from North America to South America after closing of the isthmus). The lack of difference in dispersal rate between Madagascar and the rest is plausible as well, knowing that most Malagasy dung beetles are part of one of few clades that entered Madagascar and diversified there, with very few dispersals back (Miraldo *et al*. 2011; Sole *et al*. 2011). This is also reflected by the fact that the inferred rate between Madagascar and the rest is comparatively low (Table 4). All in all, those results would support the hypothesis that access to new areas was associated with a rise of diversification rates in dung beetles. However, more fine-scale analyses are needed to confirm the implied scenarios behind this. Also, while overall sampling frequency in this phylogeny was considered in this analysis, it cannot be ruled out that some of these results are artefacts of uneven sampling between groups within the subfamily. Until an even more complete tree of Scarabaeinae is available, finding ways to correct for such sampling biases would be advisable.

## Acknowledgements

We would like to express our thankfulness for helpful discussions and comments at various stages of this project to Kimberly Sheldon, Nicole Gunter, Sergei Tarasov, Nick Matzke, Jim Fordyce, Colin Sumrall, Dave Bapst, Michelle Lawing, Jeremy Beaulieu, Luna Sanchez-Reyes, Veronica Brown, Marisol Sanchez, Todd Pierson, Hailee Korotkin, Claire Winfrey, Jess Dreyer, and Maggie Mamantov.

Finally, we thank the Society of Systematic Biologists and the Systematics Association for their financial support towards this project.

## Notes

### Competing Interest Statement

The authors have declared no competing interest.

