## Supplementary Materials for "Age, Origin, and Biogeography: Unveiling the Factors Behind the Diversification of Dung Beetles"

**Table S1 GenBank Accession Numbers**
This will be a table of all GenBank Accession numbers used in the final alignments.

| Name | 16S | 18S | 28S-D2 | 28S-D3 | CAD | COI-1 | COI-2 |
|---|---|---|---|---|---|---|---|
| *Paraphytus sp.* | | KX512471.1 | KX512561.1 | KX512638.1 | | KJ867466.1 | |
| *Frankenbergerius armatus* | GQ289676.1 | KX512504.1 | KX512593.1 | KX512664.1 | KX530439.1 | | GQ290031.1 |
| *Sarophorus costatus* | AY131523.1 | KX512496.1 | GQ289814.1 | AY131712.1 | GQ289965.1 | KF956280.1 | AY131883.1 |
| *Sarophorus tuberculatus* | AY131524.1 | JN619266.1 | | AY131713.1 | | | AY131884.1 |
| *Delopleurus pullus* | GQ289700.1 | | GQ289833.1 | GQ289918.1 | GQ289976.1 | | GQ290052.1 |
| *Coptorhina nitidipennis* | GQ289683.1 | | GQ289818.1 | GQ289901.1 | | | GQ290037.1 |
| *Odontoloma pusillum* | AY131468.1 | JN619208.1 | GQ289790.1 | AY131661.1 | GQ289930.1 | | AY131839.1 |
| *Aliuscanthoniola similaris* | JN804741.1 | | JN804810.1 | KC928076.1 | | | JN804667.1 |
| *Outenikwanus tomentosus* | GQ289748.1 | | GQ289798.1 | GQ289886.1 | GQ289948.1 | | GQ290024.1 |
| *Peckolus alpinus* | GQ289731.1 | | GQ289780.1 | GQ289869.1 | GQ289953.1 | | GQ290004.1 |
| *Endroedyolus paradoxus* | GQ289702.1 | | GQ289752.1 | GQ289838.1 | GQ289921.1 | | GQ289981.1 |
| *Silvaphilus oubosiensis* | JN804748.1 | | JN804822.1 | KC928077.1 | JN804606.1 | | JN804677.1 |
| *Namakwanus davisi* | GQ289745.1 | | GQ289802.1 | GQ289883.1 | GQ289956.1 | | GQ290021.1 |
| *Byrrhidium convexum* | GQ289732.1 | | GQ289782.1 | GQ289871.1 | GQ289938.1 | | GQ290006.1 |
| *Byrrhidium namaquensis* | GQ289724.1 | | GQ289771.1 | GQ289860.1 | GQ289931.1 | | GQ289998.1 |
| *Dicranocara deschodti* | EF656672.1 | JN969180.1 | GQ289793.1 | EF656714.1 | GQ289944.1 | | DQ667017.1 |
| *Dicranocara tatasensis* | | | | | | | DQ667010.1 |
| *Dicranocara inexpectata* | | | | | | | DQ667014.1 |
| *Boletoscapter cornutus* | AY131441.1 | JN619225.1 | | AY131632.1 | | | AY131813.1 |

| | | | | | | | |
|---|---|---|---|---|---|---|---|
| *Uroxys pygmaeus* | AY131529.1 | | | EF656712.1 | | | EF656761.1 |
| *Bdelyropsis bowditchi* | EF656654.1 | JN619144.1 | | EF656696.1 | | | EF656745.1 |
| *Uroxys micros* | AY131528.1 | | | AY131717.1 | | AY144774.1 | AY131886.1 |
| *Diorygopyx tibialis* | AY847522.1 | | | | | | |
| *Diorygopyx simpliciclunis* | AY131456.1 | JN619181.1 | | KF802118.1 | | | KF801954.1 |
| *Pseudignambia sp* | | | | | | | |
| *Diorygopyx incomptus* | AY847524.1 | | | KF802119.1 | | | KF801955.1 |
| *Diorygopyx niger* | AY847523.1 | KX512476.1 | KX512566.1 | KF802106.1 | | | KF801942.1 |
| *Onthobium cookii* | | | | | | | |
| *Ignambia fasciculata* | AY131463.1 | JN619175.1 | | AY131654.1 | | | AY131834.1 |
| *Pseudonthobium fracticolloides* | AY131477.1 | JN619182.1 | | AY131669.1 | | | AY131847.1 |
| *Paronthobium simplex* | AY131473.1 | JN619173.1 | | AY131665.1 | | | AY131843.1 |
| *Anonthobium tibiale* | AY131439.1 | JN619174.1 | | AY131630.1 | | | AY131811.1 |
| *Demarziella interrupta* | AY131511.1 | JN619178.1 | | AY131700.1 | | | |
| *Demarziella mirifica* | AY131512.1 | | | AY131701.1 | | | AY131872.1 |
| *Amphistomus complanatus* | AY131436.1 | JN619228.1 | | | | | AY131808.1 |
| *Coptodactyla meridionalis* | | | | | | | MG588139.1 |
| *Coptodactyla storeyi* | AY131497.1 | | | | | | |
| *Coptodactyla glabricollis* | AY131496.1 | JN619207.1 | | AY131687.1 | | | AY131863.1 |
| *Ochicanthon punctatus* | AY131474.1 | JN619188.1 | | AY131666.1 | | | AY131844.1 |
| *Epactoides mangabeensis* | EU030497.1 | | | EU030542.1 | | | EU030586.1 |
| *Epactoides hanskii* | EU030504.1 | GQ341988.1 | | EU030549.1 | | | EU030593.1 |
| *Epactoides helenae* | EU030507.1 | GQ341989.1 | | EU030552.1 | | | EU030595.1 |
| *Epactoides mahaboi* | EU030515.1 | GQ341991.1 | | EU030560.1 | | | EU030599.1 |
| *Epactoides semiaeneus* | EU030529.1 | | | EU030573.1 | | | EU030610.1 |
| *Epactoides tiinae* | EU030533.1 | GQ341995.1 | | EU030576.1 | | | EU030614.1 |

| | | | | | | | |
|---|---|---|---|---|---|---|---|
| *Epactoides rahagai* | EU030527.1 | GQ341994.1 | | EU030570.1 | | | EU030608.1 |
| *Epactoides major* | EU030519.1 | GQ341992.1 | | DQ369542.1 | | | EU030601.1 |
| *Epactoides femoralis* | EU030501.1 | | | EU030546.1 | | | EU030589.1 |
| *Epactoides viridicollis* | DQ369609.1 | DQ369660.1 | | DQ369543.1 | | | EU030617.1 |
| *Epactoides lissus* | EU030514.1 | | | EU030557.1 | | | EU030598.1 |
| *Epactoides incertus* | EU030509.1 | GQ341990.1 | | EU030554.1 | | | EU030597.1 |
| *Epactoides perrieri* | EU030524.1 | | | EU030569.1 | | | EU030606.1 |
| *Epactoides spinicollis* | EU030532.1 | | | EU030574.1 | | KF309738.1 | EU030612.1 |
| *Epactoides frontalis* | DQ369607.1 | DQ369661.1 | | DQ369541.1 | KF309818.1 | | EU030590.1 |
| *Coproecus hemisphaericus* | AY131451.1 | JN619177.1 | | AY131641.1 | | | AY131821.1 |
| *Lepanus occidentalis* | KF801733.1 | | | KF802062.1 | | | KY784156.1 |
| *Lepanus villosus* | | | | | | | KY784174.1 |
| *Lepanus dichrous* | | | | | | | KY784155.1 |
| *Lepanus nitidus* | AY131464.1 | | | AY131655.1 | | | AY131835.1 |
| *Lepanus australis* | KF801821.1 | | | KF802149.1 | | | KF801986.1 |
| *Lepanus palumensis* | | | | | | | KY784177.1 |
| *Lepanus politus* | AY847533.1 | | | | | | KY784151.1 |
| *Lepanus pygmaeus* | | | | | | AY144789.1 | KY784172.1 |
| *Lepanus monteithi* | | | | | | | KY784157.1 |
| *Lepanus globulus* | KF801822.1 | | | KF802150.1 | | AY144776.1 | KF801987.1 |
| *Lepanus ustulatus* | | JN619265.1 | KX512567.1 | AY131656.1 | KX530432.1 | | KY784152.1 |
| *Sauvagesinella palustris* | KF801735.1 | | | KF802064.1 | | | KF801906.1 |
| *Sauvagesinella becki* | KF801737.1 | | | KF802066.1 | | | KF801899.1 |
| *Sauvagesinella monstrosa* | KF801736.1 | | | KF802065.1 | | | KF801898.1 |
| *Monoplistes curvipes* | AY131467.1 | JN619186.1 | | AY131659.1 | | | |
| *Canthonosoma macleayi* | | KX512474.1 | KX512564.1 | KX512641.1 | | | |

| | | | | | | | |
|---|---|---|---|---|---|---|---|
| *Canthonosoma castelnaui* | AY131447.1 | | | AY131638.1 | | | AY131818.1 |
| *Saphobius squamulosus* | AY131480.1 | | | AY131672.1 | | | |
| *Saphobius setosus* | AY131479.1 | JN619310.1 | | AY131671.1 | | | |
| *Cephalodesmius armiger* | AY131448.1 | AY745577.1 | | | | | |
| *Cephalodesmius quadridens* | AY131449.1 | | | AY131639.1 | | | AY131819.1 |
| *Monteithocanthon glaber* | EF656646.1 | | | EF656688.1 | | AY144792.1 | KY784166.1 |
| *Temnoplectron boucomonti* | | | | | | AY144791.1 | AY144792.1 |
| *Temnoplectron major* | | | | | | AY144788.1 | AY144791.1 |
| *Temnoplectron disruptum* | | | | | | AY144785.1 | AY144788.1 |
| *Temnoplectron cooki* | | | | | | AY144787.1 | AY144785.1 |
| *Temnoplectron reyi* | | | | | | AY144786.1 | AY144787.1 |
| *Temnoplectron politulum* | AY131484.1 | | | AY131676.1 | | AY144782.1 | AY144786.1 |
| *Temnoplectron aeneopiceum* | | | | | | AY144783.1 | AY144782.1 |
| *Temnoplectron subvolitans* | | | | | | AY144779.1 | AY144783.1 |
| *Temnoplectron monteithi* | | | | | | AY144790.1 | AY144794.1 |
| *Temnoplectron finnigani* | AY131483.1 | JN619220.1 | | AY131675.1 | | AY144771.1 | AY131851.1 |
| *Temnoplectron lewisense* | | | | | | | AY144795.1 |
| *Janssensantus sp* | | | | | | | |
| *Tanzanolus sp* | | | | | | | |
| *Gyronotus pumilus* | GQ289714.1 | GQ342009.1 | GQ289765.1 | GQ289851.1 | GQ289950.1 | | GQ342148.1 |
| *Canthodimorpha lawrencei* | GQ289744.1 | | GQ289797.1 | GQ289889.1 | | | GQ290018.1 |
| *Anachalcos convexus* | AY131437.1 | GQ341942.1 | GQ289768.1 | AY131628.1 | GQ289929.1 | | AY131809.1 |
| *Anachalcos suturalis* | AY131438.1 | | | AY131629.1 | | | AY131810.1 |
| *Paragymnopleurus maurus* | AY131545.1 | JN619216.1 | | AY131733.1 | | | AY131902.1 |
| *Paragymnopleurus striatus* | AY131546.1 | | | AY131734.1 | | | AY131903.1 |
| *Allogymnopleurus thalassinus* | JN804680.1 | AY821521.1 | JN804751.1 | AY131729.1 | JN804566.1 | | AY131898.1 |

| | | | | | | | |
|---|---|---|---|---|---|---|---|
| *Garreta nitens* | AY131542.1 | | JN804773.1 | AY131730.1 | JN804579.1 | | AY131899.1 |
| *Gymnopleurus virens* | AY131543.1 | JN619219.1 | | AY131731.1 | | | AY131900.1 |
| *Gymnopleurus humanus* | JN804700.1 | | JN804772.1 | | | | JN804627.1 |
| *Gymnopleurus flagellatus* | | | | | | KT454096.1 | AY039364.1 |
| *Gymnopleurus mopsus* | KJ721833.1 | | | KJ721895.1 | | DQ369595.1 | AY039363.1 |
| *Arachnodes kelifelyi* | DQ369620.1 | DQ369673.1 | | DQ369554.1 | | | GQ342120.1 |
| *Arachnodes grossepunctatus* | GQ341890.1 | | | GQ342052.1 | | | GQ342119.1 |
| *Arachnodes pusillus* | GQ341901.1 | GQ341970.1 | | GQ342064.1 | | | GQ342126.1 |
| *Arachnodes philippi* | GQ341899.1 | GQ341968.1 | | GQ342062.1 | | | |
| *Arachnodes micheli* | GQ341925.1 | GQ342004.1 | | GQ342091.1 | | DQ369590.1 | GQ342143.1 |
| *Arachnodes andriai* | DQ369614.1 | DQ369666.1 | | DQ369548.1 | | DQ369598.1 | GQ342115.1 |
| *Arachnodes manaitrai* | DQ369625.1 | GQ341964.1 | | DQ369559.1 | | | GQ342124.1 |
| *Pseudoarachnodes mantillerii* | GQ341937.1 | | | GQ342107.1 | | DQ369600.1 | |
| *Pseudoarachnodes hanskii* | GQ341935.1 | DQ369678.1 | | GQ342105.1 | | | GQ342155.1 |
| *Pseudoarachnodes semichalceus* | GQ341938.1 | | | GQ342108.1 | | | GQ342156.1 |
| *Pseudoarachnodes insularis* | GQ341936.1 | GQ342029.1 | | GQ342106.1 | | | |
| *Arachnodes genieri* | GQ341923.1 | GQ342002.1 | | GQ342089.1 | | DQ369591.1 | GQ342141.1 |
| *Arachnodes apotolamproides* | DQ369616.1 | DQ369669.1 | KX512574.1 | DQ369550.1 | | | GQ342136.1 |
| *Arachnodes antoetrae* | DQ369615.1 | DQ369667.1 | | DQ369549.1 | | | EU248061.1 |
| *Arachnodes morio* | GQ341897.1 | GQ341967.1 | | GQ342060.1 | | | GQ342125.1 |
| *Arachnodes colasi* | | GQ341957.1 | | | | DQ369597.1 | GQ342116.1 |
| *Arachnodes sicardi* | DQ369623.1 | DQ369677.1 | | DQ369557.1 | | | GQ342130.1 |
| *Arachnodes emmae* | GQ341920.1 | GQ342001.1 | | GQ342086.1 | | | GQ342138.1 |
| *Arachnodes globuloides* | GQ341889.1 | | | GQ342051.1 | | | GQ342118.1 |
| *Arachnodes seminitidus* | GQ341904.1 | GQ341974.1 | | GQ342068.1 | | | GQ342129.1 |
| *Arachnodes delphinensis* | GQ341919.1 | GQ342000.1 | | GQ342085.1 | | | EU248060.1 |

| | | | | | | | |
|---|---|---|---|---|---|---|---|
| *Arachnodes splendidus* | EF656678.1 | GQ342007.1 | | EF656720.1 | | | GQ342146.1 |
| *Arachnodes prasinus* | GQ341926.1 | GQ342005.1 | | GQ342092.1 | | | GQ342144.1 |
| *Arachnodes ruteri* | GQ341927.1 | | | GQ342093.1 | | DQ369593.1 | GQ342145.1 |
| *Arachnodes cuprarius* | DQ369618.1 | DQ369671.1 | | DQ369552.1 | | | DQ369478.1 |
| *Arachnodes mantasoae* | GQ341924.1 | GQ342003.1 | | GQ342090.1 | | | GQ342142.1 |
| *Arachnodes mahafalyensis* | GQ341893.1 | GQ341963.1 | | GQ342056.1 | | | GQ342123.1 |
| *Arachnodes saprinoides* | DQ369621.1 | DQ369675.1 | | DQ369555.1 | | | GQ342128.1 |
| *Arachnodes purpuricollis* | GQ341900.1 | GQ341969.1 | | GQ342063.1 | | | |
| *Arachnodes luctuosus* | | | | GQ342054.1 | | | GQ342121.1 |
| *Arachnodes robinsoni* | | GQ341971.1 | | GQ342065.1 | | | GQ342127.1 |
| *Arachnodes dichrous* | DQ369624.1 | DQ369668.1 | | DQ369558.1 | | | |
| *Litocopris muticus* | JN804711.1 | | JN804782.1 | | JN804585.1 | | JN804639.1 |
| *Copris jacchoides* | | | | | | | JN804618.1 |
| *Copris hispanicus* | | | | | | | AY039366.1 |
| *Copris sonensis* | | | | | | KM441825.1 | MG642091.1 |
| *Copris lunaris* | | | | | | | AY039365.1 |
| *Copris agnus* | AY131490.1 | | | AY131682.1 | | | AY131857.1 |
| *Copris laeviceps* | AY131492.1 | | | | | | |
| *Paracopris punctulatus* | | EF188042.1 | | | | | EF188219.1 |
| *Copris lugubris* | AY131493.1 | AY821529.1 | | AY131684.1 | | | AY131860.1 |
| *Copris aeneus* | AY131489.1 | | | AY131681.1 | | | AY131856.1 |
| *Copris cornifrons* | JN804686.1 | | JN804759.1 | | | | JN804613.1 |
| *Copris amyntor* | AY131491.1 | | DQ430882.1 | AY131683.1 | | | AY131858.1 |
| *Copris confucius* | | EF187978.1 | | EF188048.1 | | | EF188135.1 |
| *Copris sinicus* | AY131495.1 | | | AY131686.1 | | | AY131862.1 |
| *Deltochilum carinatum* | AY131453.1 | | | AY131644.1 | | | AY131824.1 |

| | | | | | | | |
|---|---|---|---|---|---|---|---|
| *Deltochilum barbipes* | | | | AY131643.1 | | | AY131823.1 |
| *Deltochilum pseudoparile* | AY131455.1 | | | AY131646.1 | | | AY131826.1 |
| *Deltochilum mexicanum* | | KX512461.1 | DQ430879.1 | KX512630.1 | | | |
| *Deltochilum amazonicum* | AY131452.1 | | | AY131642.1 | | | AY131822.1 |
| *Deltochilum gibbosum* | AY131454.1 | DQ012274.1 | DQ430878.1 | DQ430930.1 | | | AY131825.1 |
| *Eudinopus dytiscoides* | AY131461.1 | DQ430832.1 | DQ430880.1 | AY131652.1 | | | AY131832.1 |
| *Hansreia affinis* | AY131462.1 | JN619191.1 | | AY131653.1 | | | AY131833.1 |
| *Megathoposoma candezei* | AY131465.1 | JN619142.1 | | AY131657.1 | | | AY131836.1 |
| *Megathopa villosa* | | | DQ430875.1 | DQ430927.1 | | | |
| *Malagoniella puncticollis* | | DQ430830.1 | DQ430874.1 | DQ430926.1 | | | |
| *Scybalophagus plicatipennis* | | KX512440.1 | DQ430876.1 | DQ430928.1 | KX530423.1 | | |
| *Scybalophagus rugosus* | | | DQ430877.1 | DQ430929.1 | | | |
| *Canthon cyanellus* | KX807615.1 | | | | | | KX807648.1 |
| *Canthon lamprimus* | EF656648.1 | | | EF656690.1 | | | EF656739.1 |
| *Canthon perseverans* | GQ341916.1 | GQ341985.1 | | GQ342080.1 | | | |
| *Scybalocanthon pygidialis* | AY131481.1 | JN619189.1 | | AY131673.1 | | | AY131849.1 |
| *Canthon deyrollei* | GQ341915.1 | GQ341984.1 | | GQ342078.1 | | | |
| *Canthon indigaceus* | AY131443.1 | | | AY131634.1 | | | AY131814.1 |
| *Canthon aequinoctialis* | GQ341914.1 | GQ341983.1 | KX512553.1 | KX512631.1 | | | |
| *Canthon luteicollis* | AY131444.1 | | | AY131635.1 | | | AY131815.1 |
| *Canthon viridis* | AY131446.1 | AY821531.1 | | AY131637.1 | | MG321659.1 | AY131817.1 |
| *Phanaeus sororibispinus* | | | | | | EU477307.1 | MG321647.1 |
| *Phanaeus alvarengai* | | | EU432232.1 | | | EU477294.1 | |
| *Phanaeus haroldi* | | | EU432221.1 | | | EU477296.1 | |
| *Phanaeus melibaeus* | | | EU432224.1 | | | EU477309.1 | |
| *Phanaeus kirbyi* | | | EU432234.1 | | | EU477313.1 | |

| | | | | | | | |
|---|---|---|---|---|---|---|---|
| *Phanaeus paleano* | | | EU432235.1 | | | MG321655.1 | |
| *Oxysternon conspicillatum* | AY131608.1 | JN619205.1 | MG321631.1 | AY131792.1 | | EU477358.1 | AY131948.1 |
| *Oxysternon durantoni* | | | EU432273.1 | | | MG321670.1 | |
| *Oxysternon macleayi* | | | | | | EU477357.1 | |
| *Oxysternon festivum* | | | EU432272.1 | | | EU477291.1 | |
| *Phanaeus splendidulus* | | | EU432222.1 | | | EU477292.1 | |
| *Phanaeus dejeani* | | | EU432223.1 | | | EU477362.1 | |
| *Oxysternon silenus* | | | EU432271.1 | | | MG321656.1 | |
| *Diabroctis mirabilis* | | | MG321632.1 | | | | MG321645.1 |
| *Sulcophanaeus batesi* | | | DQ430856.1 | DQ430910.1 | | MG321673.1 | |
| *Coprophanaeus bellicosus* | | | | | | MG321651.1 | MG321638.1 |
| *Coprophanaeus bonariensis* | | | MG321627.1 | | | MG321667.1 | MG321640.1 |
| *Coprophanaeus lancifer* | AY131604.1 | | MG321633.1 | AY131788.1 | | MG321650.1 | AY131945.1 |
| *Coprophanaeus ensifer* | | | MG321624.1 | | | | MG321641.1 |
| *Dendropaemon bahianus* | | | | | | | |
| *Coprophanaeus chiriquensis* | | KX512455.1 | KX512545.1 | KX512628.1 | KX530427.1 | EU477349.1 | |
| *Coprophanaeus pluto* | | | EU432265.1 | | | EU477353.1 | |
| *Coprophanaeus ignecinctus* | | | EU432267.1 | | | MG321671.1 | |
| *Coprophanaeus telamon* | AY131605.1 | JN619140.1 | | AY131789.1 | | MG321663.1 | AY131946.1 |
| *Coprophanaeus dardanus* | | | | | | EU477300.1 | |
| *Phanaeus achilles* | | | EU432228.1 | | | EU477321.1 | |
| *Phanaeus prasinus* | | | EU432240.1 | | | EU477299.1 | |
| *Phanaeus chalcomelas* | | | EU432226.1 | | | EU477306.1 | |
| *Phanaeus lecourti* | | | EU432231.1 | | | EU477304.1 | |
| *Phanaeus meleagris* | | | EU432230.1 | | | | |
| *Phanaeus demon* | AY131610.1 | | | | | | AY131950.1 |

| | | | | | | | |
|---|---|---|---|---|---|---|---|
| *Phanaeus amethystinus amethystinus* | | | | | | EU477329.1 | |
| *Phanaeus lunaris* | | | EU432249.1 | | | EU477337.1 | |
| *Phanaeus howdeni* | | | EU432255.1 | | | EU477338.1 | |
| *Phanaeus sallei* | AY131611.1 | AY821524.1 | EU432256.1 | AY131793.1 | | EU477333.1 | AY131951.1 |
| *Phanaeus amithaon* | | | EU432253.1 | | | | |
| *Phanaeus wagneri wagneri* | | | | | | EU477319.1 | |
| *Phanaeus pyrois* | | | EU432239.1 | | | EU477317.1 | |
| *Phanaeus endymion* | | | EU432237.1 | | | EU477345.1 | |
| *Phanaeus igneus* | | DQ430828.1 | DQ430855.1 | DQ430909.1 | | EU477347.1 | |
| *Phanaeus vindex* | | | EU432264.1 | | | | |
| *Phanaeus triangularis texensis* | | | | | | EU477341.1 | |
| *Phanaeus quadridens* | | | EU432259.1 | | | EU477336.1 | |
| *Phanaeus yecoraensis* | | | EU432254.1 | | | EU477323.1 | |
| *Phanaeus nimrod* | | | EU432241.1 | | | EU477324.1 | |
| *Phanaeus furiosus* | | | EU432244.1 | | | | |
| *Ennearabdus lobocephalus* | AY131532.1 | DQ430829.1 | DQ430861.1 | AY131721.1 | | | AY131889.1 |
| *Eucranium arachnoides* | AY131533.1 | AY821527.1 | DQ430859.1 | AY131722.1 | | | AY131890.1 |
| *Eucranium planicolle* | | | DQ430860.1 | DQ430913.1 | | | |
| *Anomiopsoides cavifrons* | | | DQ430865.1 | DQ430918.1 | | | |
| *Glyphoderus sterquilinus* | AY131534.1 | | DQ430863.1 | AY131723.1 | | | AY131891.1 |
| *Anomiopsoides biloba* | AY131530.1 | | DQ430864.1 | AY131719.1 | | | AY131887.1 |
| *Glyphoderus centralis* | | | DQ430862.1 | DQ430915.1 | | | |
| *Anomiopsoides heteroclyta* | AF499693.1 | JN619203.1 | DQ430866.1 | AY131720.1 | | | AY131888.1 |
| *Canthidium haroldi* | AY131506.1 | | | AY131695.1 | | | AY131868.1 |
| *Kheper subaeneus* | AF499696.1 | | | | | | |
| *Kheper nigroaeneus* | AF499695.1 | AY821523.1 | DQ430870.1 | AY131795.1 | | | AY131953.1 |

| | | | | | | | |
|---|---|---|---|---|---|---|---|
| *Kheper bonellii* | JN804740.1 | | JN982317.1 | | JN819271.1 | AY258257.1 | JN804665.1 |
| *Pachysoma valeflorae* | | | | | | AY258247.1 | AY258257.1 |
| *Pachysoma schinzi* | | | | | | AY258213.1 | AY258247.1 |
| *Pachysoma aesculapius* | | | | | | AY258223.1 | AY258213.1 |
| *Pachysoma endroeydi* | | | | | | AY258226.1 | AY258223.1 |
| *Pachysoma glentoni* | | | | | | AY258215.1 | AY258226.1 |
| *Pachysoma hippocrates* | AF499699.1 | | DQ430868.1 | DQ430920.1 | | AY258253.1 | AY258215.1 |
| *Pachysoma denticollis* | | | | | | AY258241.1 | AY258253.1 |
| *Pachysoma rotundigenus* | | | | | | AY258244.1 | AY258241.1 |
| *Pachysoma rodriguesi* | | | | | | AY258250.1 | AY258244.1 |
| *Pachysoma striatus* | | | | | | AY258231.1 | AY258250.1 |
| *Pachysoma gariepinus* | | | | | | AY258238.1 | AY258231.1 |
| *Pachysoma bennigseni* | AF499698.1 | | | | | | AY258238.1 |
| *Scarabaeus cicatricosus* | | | | | | | AY039362.1 |
| *Drepanopodus costatus* | AY131612.1 | JN619206.1 | | AY131794.1 | | | AY131952.1 |
| *Scarabaeus galenus* | AF499704.1 | | | AY131798.1 | | AY965239.1 | AY131956.1 |
| *Drepanopodus proximus* | AF499694.1 | | | | | | AY965239.1 |
| *Scarabaeus bohemani* | AF499701.1 | | | | | | |
| *Scarabaeus flavicornis* | AF499702.1 | | | | | | |
| *Scarabaeus brittoni* | AY131618.1 | JN619308.1 | | AY131800.1 | | | AY131958.1 |
| *Scarabaeus adamastor* | AF499711.1 | | | | | | |
| *Scarabaeus hippias* | AY131619.1 | | | AY131801.1 | | | AY131959.1 |
| *Pachylomera femoralis* | | | | | | | |
| *Scarabaeus rugosus* | AF499706.1 | | | | | | FJ763730.1 |
| *Scarabaeus caffer* | JN804737.1 | | JN804806.1 | | JN819270.1 | | |
| *Scarabaeus westwoodi* | AF499709.1 | | | | | GU305943.1 | |

| | | | | | | | |
|---|---|---|---|---|---|---|---|
| *Scarabaeus viettei* | GU305941.1 | | | | | | GU305943.1 |
| *Scarabaeus goryi* | AF499705.1 | | | | | | |
| *Scarabaeus satyrus* | AF499708.1 | | | | | GU305942.1 | |
| *Scarabaeus radama* | GU305940.1 | | | | | KP752076.1 | GU305942.1 |
| *Scarabaeus sacer* | | | | | | KP752063.1 | KP752076.1 |
| *Scarabaeus pius* | | | | | | KP752070.1 | KP752055.1 |
| *Scarabaeus thphon* | | | | | | | KP752068.1 |
| *Neateuchus proboscideus* | AF499697.1 | | | | | | |
| *Scarabaeus deludens* | | KP419281.1 | DQ430869.1 | GU226585.1 | | | |
| *Scarabaeus zambezianus* | AF499710.1 | | JN804808.1 | | JN819268.1 | | |
| *Leotrichillum sp* | | | | | | | |
| *Trichillidium pilosum* | | KX512459.1 | KX512549.1 | | | | |
| *Canthidium guanacaste* | AY131505.1 | AY821530.1 | | AY131694.1 | | | AY131867.1 |
| *Canthidium rufinum* | AY131507.1 | | | AY131696.1 | | | AY131869.1 |
| *Canthidium thalassinum* | AY131508.1 | | | AY131697.1 | | | AY131870.1 |
| *Dichotomius boreus* | AY131514.1 | | | AY131703.1 | | HQ824542.1 | AY131874.1 |
| *Dichotomius laevicollis* | HQ824536.1 | | | | | | |
| *Dichotomius yucatanus* | AY131516.1 | | DQ430850.1 | AY131705.1 | | | AY131876.1 |
| *Dichotomius parcepunctatus* | AY131515.1 | | | AY131704.1 | | HQ824539.1 | AY131875.1 |
| *Dichotomius geminatus* | HQ824533.1 | | | | | HQ824541.1 | |
| *Dichotomius semisquamosus* | HQ824535.1 | | | | | HQ824540.1 | |
| *Dichotomius sericeus* | HQ824534.1 | | | | | | |
| *Dichotomius nisus* | HQ824538.1 | | | | | HQ824543.1 | |
| *Dichotomius bos* | HQ824537.1 | | | | | | |
| *Ateuchus viduus* | | | | | | | |
| *Ateuchus ecuadorense* | EF656650.1 | | | EF656692.1 | | | EF656741.1 |

| | | | | | | | |
|---|---|---|---|---|---|---|---|
| *Ateuchus chrysopyge* | AY131502.1 | JN619143.1 | | AY131692.1 | | | AY131866.1 |
| *Ateuchus floridensis* | | | | | | | |
| *Eurysternus inflexus* | AY131538.1 | | | AY131726.1 | | | AY131895.1 |
| *Eurysternus plebejus* | AY131539.1 | | | AY131727.1 | | | AY131896.1 |
| *Eurysternus angustulus* | AY131535.1 | AY821533.1 | | AY131724.1 | | | AY131892.1 |
| *Eurysternus velutinus* | AY131540.1 | | | AY131728.1 | | | AY131897.1 |
| *Eurysternus caribaeus* | AY131536.1 | | | AY131725.1 | | | AY131893.1 |
| *Eurysternus hamaticollis* | AY131537.1 | | | EF656708.1 | | KF309775.1 | AY131894.1 |
| *Nanos minutus* | EU247965.1 | GQ342019.1 | | EU248016.1 | KF309836.1 | KF309746.1 | EU248068.1 |
| *Nanos bicoloratus* | EU247994.1 | | | EU248047.1 | KF309819.1 | KF309755.1 | EU248089.1 |
| *Nanos humeralis* | | | | KF309915.1 | KF309825.1 | KF309777.1 | |
| *Nanos rubrosignatus* | | GQ342022.1 | | EU248013.1 | KF309838.1 | KF309753.1 | EU248065.1 |
| *Nanos hanskii* | DQ369632.1 | DQ369679.1 | | DQ369566.1 | KF309824.1 | KF309792.1 | EU248074.1 |
| *Nanos nitens* | EU247989.1 | | | EU248041.1 | KF309829.1 | DQ369604.1 | EU248182.1 |
| *Nanos viettei* | DQ369631.1 | DQ369684.1 | | DQ369565.1 | KF309842.1 | KF309751.1 | EU248153.1 |
| *Nanos clypeatus* | EU247970.1 | GQ342013.1 | | EF656718.1 | KF309822.1 | KF309780.1 | EU248106.1 |
| *Nanos dubitatus* | DQ369629.1 | DQ369682.1 | | DQ369563.1 | KF309834.1 | KF309767.1 | EU248099.1 |
| *Nanos occidentalis* | DQ369633.1 | DQ369680.1 | | DQ369567.1 | KF309830.1 | KF309769.1 | EU248087.1 |
| *Nanos peyrierasi* | DQ369630.1 | DQ369683.1 | | DQ369564.1 | KF743767.1 | KF309782.1 | EU248092.1 |
| *Nanos vadoni* | EU248003.1 | GQ342024.1 | | EU248054.1 | KF309841.1 | KF309759.1 | EU248097.1 |
| *Nanos manomboensis* | EU247981.1 | GQ342015.1 | | EU248033.1 | KF743770.1 | KF309747.1 | EU248078.1 |
| *Nanos bimaculatus* | DQ369628.1 | DQ369681.1 | | DQ369562.1 | KF309820.1 | KF309750.1 | EU248071.1 |
| *Nanos binotatus* | | | | | KF309821.1 | DQ369596.1 | HM029151.1 |
| *Arachnodes semipunctatus* | DQ369622.1 | DQ369676.1 | | DQ369556.1 | | KF309739.1 | DQ369481.1 |
| *Nanos semiscribosus* | GQ341903.1 | GQ341973.1 | | KF309931.1 | KF309839.1 | KF309703.1 | GQ342150.1 |
| *Apotolamprus cyanescens* | GQ341886.1 | GQ341958.1 | | KF309887.1 | KF309803.1 | KF309707.1 | GQ342111.1 |

| | | | | | | | |
|---|---|---|---|---|---|---|---|
| *Apotolamprus latipennis* | GQ341873.1 | GQ341946.1 | | GQ342034.1 | KF309807.1 | KF309733.1 | GQ342113.1 |
| *Apotolamprus zombitsyensis* | GQ341879.1 | GQ341950.1 | | KF309904.1 | KF309816.1 | KF309710.1 | |
| *Apotolamprus metallicus* | | | | KF309894.1 | KF309810.1 | DQ369592.1 | |
| *Arachnodes balianus* | DQ369617.1 | DQ369670.1 | | DQ369551.1 | | KF309726.1 | DQ369477.1 |
| *Apotolamprus milloti* | GQ341876.1 | | | KF309895.1 | KF309811.1 | KF309716.1 | |
| *Apotolamprus vadoni* | | | | KF309902.1 | | KF309702.1 | |
| *Apotolamprus ambohitsitondronensis* | | | | KF309885.1 | KF309802.1 | KF309709.1 | |
| *Apotolamprus marojejyensis* | GQ341874.1 | | | GQ342035.1 | KF309809.1 | KF309728.1 | |
| *Apotolamprus quadrimaculatus* | GQ341877.1 | | | KF309899.1 | KF309813.1 | KF309715.1 | |
| *Apotolamprus sericeus* | | | | KF309901.1 | KF309815.1 | KF309706.1 | |
| *Apotolamprus helenae* | DQ369611.1 | DQ369663.1 | | DQ369545.1 | KF309806.1 | KF309705.1 | EU248063.1 |
| *Apotolamprus hanskii* | DQ369610.1 | DQ369662.1 | | DQ369544.1 | KF309805.1 | KF309714.1 | EU248062.1 |
| *Apotolamprus quadrinotatus* | DQ369612.1 | DQ369664.1 | | DQ369546.1 | KF309814.1 | KF309704.1 | EU030619.1 |
| *Apotolamprus darainaensis* | GQ341872.1 | GQ341943.1 | | KF309888.1 | KF309804.1 | KF309712.1 | GQ342112.1 |
| *Apotolamprus orangeaensis* | | | | KF309896.1 | | KF309713.1 | |
| *Apotolamprus pseudomanomboensis* | | | | KF309898.1 | KF309812.1 | KF309708.1 | |
| *Apotolamprus manomboensis* | GQ341895.1 | GQ341965.1 | | GQ342058.1 | KF309808.1 | KF309779.1 | GQ342114.1 |
| *Apotolamprus sahatezaensis* | | | | KF309932.1 | KF309840.1 | KF309737.1 | |
| *Cambefortantus ranomafanensis* | GQ341913.1 | GQ341982.1 | | KF309905.1 | KF309817.1 | | |
| *Bohepilissus subtilis* | | | GQ289784.1 | GQ289873.1 | GQ289940.1 | | GQ290008.1 |
| *Heliocopris andersoni* | AY131518.1 | AY821526.1 | | AY131707.1 | | | AY131878.1 |
| *Heliocopris hamadryas* | AY131519.1 | | GQ289827.1 | AY131708.1 | GQ289971.1 | | AY131879.1 |
| *Metacatharsius opacus* | AY131498.1 | JN619202.1 | | AY131688.1 | | | AY131864.1 |
| *Metacatharsius exiguiformis* | JN804720.1 | | JN804788.1 | | JN804589.1 | | JN804647.1 |
| *Metacatharsius marani* | JN804721.1 | | JN804791.1 | | JN804591.1 | | JN804650.1 |
| *Catharsius philus* | AY131487.1 | | | AY131679.1 | | | AY131854.1 |

| | | | | | | | |
|---|---|---|---|---|---|---|---|
| *Catharsius calaharicus* | AY131485.1 | JN619192.1 | | AY131677.1 | | | AY131852.1 |
| *Catharsius sesostris* | AY131488.1 | | | AY131680.1 | | JQ855856.1 | AY131855.1 |
| *Catharsius molossus* | AY131486.1 | | | KJ721870.1 | | | AY131853.1 |
| *Epirinus aeneus* | AY131458.1 | JN619201.1 | GQ289777.1 | AY131649.1 | GQ289935.1 | | AY131829.1 |
| *Epirinus hilaris* | AY131459.1 | | | AY131650.1 | | | AY131830.1 |
| *Epirinus convexus* | GQ289705.1 | | GQ289755.1 | GQ289841.1 | GQ289924.1 | | GQ289984.1 |
| *Epirinus silvestris* | GQ289750.1 | | GQ289800.1 | GQ289882.1 | GQ289946.1 | | GQ290019.1 |
| *Epirinus aquilus* | | | HQ289943.1 | | HQ289923.1 | | HQ289975.1 |
| *Epirinus sebastiani* | HQ289917.1 | | HQ289965.1 | | HQ289939.1 | | HQ289976.1 |
| *Epirinus relictus* | HQ289915.1 | | HQ289964.1 | | HQ289937.1 | | HQ290003.1 |
| *Epirinus ngomae* | HQ289909.1 | | HQ289957.1 | | HQ289933.1 | | HQ289987.1 |
| *Epirinus hluhluwensis* | HQ289906.1 | | HQ289955.1 | | HQ289931.1 | | HQ289995.1 |
| *Epirinus comosus* | HQ289899.1 | | HQ289946.1 | | HQ289926.1 | | HQ289991.1 |
| *Epirinus pygidialis* | | | HQ289963.1 | | | | HQ289994.1 |
| *Epirinus obtusus* | HQ289912.1 | | HQ289962.1 | | HQ289935.1 | | HQ289982.1 |
| *Epirinus scrobiculatus* | HQ289902.1 | | HQ289948.1 | | HQ289927.1 | | HQ289979.1 |
| *Epirinus sulcipennis* | HQ289918.1 | | HQ289969.1 | | HQ289941.1 | | HQ289988.1 |
| *Epirinus flagellatus* | HQ289903.1 | | HQ289951.1 | | HQ289928.1 | | HQ289983.1 |
| *Epirinus validus* | HQ289920.1 | | HQ289971.1 | | | | HQ289984.1 |
| *Onitis caffer* | AY131600.1 | | JN804798.1 | EF656713.1 | | | EF656762.1 |
| *Onitis ion* | | | | | | | AY039340.1 |
| *Bubas bison* | AY131595.1 | | | AY131779.1 | | | AY131938.1 |
| *Bubas bubalus* | AY131596.1 | | | AY131780.1 | | | AY131939.1 |
| *Cheironitis scabrosus* | JN804690.1 | | JN804763.1 | | JN804573.1 | | JN804616.1 |
| *Cheironitis hoplosternus* | AY131597.1 | AY821528.1 | | AY131781.1 | | | AY131940.1 |
| *Onitis alexis* | AY131599.1 | DQ430835.1 | DQ430888.1 | AY131783.1 | | | AY131942.1 |

| | | | | | | | |
|---|---|---|---|---|---|---|---|
| *Onitis falcatus* | AY131601.1 | | | AY131785.1 | | | AY131943.1 |
| *Onitis subopacus* | | EF188030.1 | | | | | EF188205.1 |
| *Hammondantus psammophilus* | GQ289743.1 | KX512501.1 | GQ289796.1 | GQ289881.1 | | | GQ290017.1 |
| *Pycnopanelus krikkeni* | GQ289708.1 | | GQ289761.1 | GQ289845.1 | | | GQ289987.1 |
| *Xinidium dentilabris* | GQ289670.1 | KX512493.1 | GQ289805.1 | GQ289890.1 | GQ289958.1 | | GQ290025.1 |
| *Xinidium dewitzi* | GQ289690.1 | | GQ289824.1 | GQ289908.1 | GQ289969.1 | | GQ290042.1 |
| *Macroderes amplior* | GQ289698.1 | | GQ289834.1 | GQ289916.1 | GQ289978.1 | | GQ290050.1 |
| *Macroderes minutus* | GQ289699.1 | | GQ289835.1 | GQ289915.1 | GQ289977.1 | | GQ290051.1 |
| *Macroderes mutilans* | GQ289673.1 | | GQ289808.1 | GQ289893.1 | GQ289959.1 | | GQ290028.1 |
| *Macroderes mutilatus* | | | | | | | |
| *Dwesasilvasedis medinae* | GQ289711.1 | KX512508.1 | GQ289762.1 | GQ289848.1 | GQ289926.1 | | GQ289990.1 |
| *Sisyphus fasciculatus* | | | | | | | |
| *Sisyphus crispatus* | AY131624.1 | | | AY131805.1 | | KM452277.1 | AY131963.1 |
| *Sisyphus schaefferi* | KJ721831.1 | | | KJ721893.1 | | | AY039367.1 |
| *Sisyphus seminulum* | AY131627.1 | | | | | | AY131966.1 |
| *Neosisyphus fortuitus* | AY131621.1 | AY821534.1 | | AY131803.1 | | | AY131961.1 |
| *Sisyphus gazanus* | AY131626.1 | | | AY131807.1 | | | AY131965.1 |
| *Phalops rufosignatus* | JN804732.1 | | JN804801.1 | | JN804601.1 | | JN804660.1 |
| *Phalops ardea* | AY131592.1 | JN619194.1 | | AY131776.1 | | | AY131935.1 |
| *Onthophagus signatus* | JN804727.1 | EF188038.1 | | EF188124.1 | | EU162450.1 | EF188215.1 |
| *Digitonthophagus gazella* | AY131563.1 | EF188036.1 | DQ430884.1 | FJ817954.1 | | | AY131918.1 |
| *Proagoderus bicallosus* | | | | | | | |
| *Proagoderus schwaneri* | AY131594.1 | | | AY131778.1 | | | AY131937.1 |
| *Parascatonomus penicillatus* | DQ369540.1 | DQ369585.1 | | DQ369522.1 | | | EF188221.1 |
| *Onthophagus semiareus* | AY131589.1 | | | AY131773.1 | | | AY131932.1 |
| *Onthophagus seniculus* | | | | | | | |

| | | | | | | | |
|---|---|---|---|---|---|---|---|
| *Serrophorus seniculus* | | EF188045.1 | | EF188131.1 | | | EF188225.1 |
| *Onthophagus diabolicus* | | | | | | | |
| *Onthophagus avocetta* | | EF188031.1 | | EF188115.1 | | | |
| *Onthophagus elegans* | | EF188033.1 | | EF188117.1 | | | EF188208.1 |
| *Proagoderus aciculatus* | JN804735.1 | | JN804804.1 | | JN804602.1 | EU162465.1 | JN804663.1 |
| *Onthophagus rangifer* | EU162562.1 | | | | | | |
| *Onthophagus lanista* | EU162551.1 | | | | | | |
| *Onthophagus tersidorsis* | EU162574.1 | | | | | | |
| *Oniticellus planatus* | | EF188028.1 | | EF188113.1 | | | EF188203.1 |
| *Liatongus vertagus* | | EF188025.1 | | EF188110.1 | | | EF188201.1 |
| *Tiniocellus spinipes* | AY131556.1 | EF188046.1 | JN804821.1 | AY131743.1 | | | AY131912.1 |
| *Proagoderus sapphirinus* | | | | | | | |
| *Tragiscus dimidiatus* | AY131557.1 | EF188047.1 | | AY131744.1 | | | AY131913.1 |
| *Oniticellus egregius* | AY131553.1 | | JN804795.1 | AY131740.1 | JN804595.1 | | AY131909.1 |
| *Drepanocerus orientalis* | | | | | | | |
| *Drepanocerus kirbyi* | AY131549.1 | | | AY131737.1 | | | AY131906.1 |
| *Drepanocerus laticollis* | | EF187980.1 | | EF188049.1 | | | EF188137.1 |
| *Drepanocerus bechynei* | AY131548.1 | JN619217.1 | | AY131736.1 | | | AY131905.1 |
| *Eodrepanus bechynei* | | | | | | KM439695.1 | |
| *Euoniticellus fulvus* | AY131554.1 | AY821522.1 | | AY131741.1 | | | AY131910.1 |
| *Euoniticellus triangulatus* | | EF187983.1 | | EF188050.1 | | | EF188138.1 |
| *Euoniticellus intermedius* | AY131550.1 | JN619209.1 | KX512609.1 | AY131738.1 | | | |
| *Euoniticellus africanus* | JN804699.1 | | JN804771.1 | | JN804578.1 | | JN804626.1 |
| *Tiniocellus sarawacus* | AY131555.1 | JN619222.1 | | AY131742.1 | | | AY131911.1 |
| *Liatongus militaris* | AY131552.1 | EF188024.1 | JN804783.1 | AY131739.1 | JN804587.1 | | EF188199.1 |
| *Liatongus phanaeoides* | KJ721841.1 | | | KJ721903.1 | | | |

| | | | | | | | |
|---|---|---|---|---|---|---|---|
| *Cyptochirus ambiguus* | AY131547.1 | | JN804766.1 | AY131735.1 | JN804574.1 | | AY131904.1 |
| *Helictopleurus fungicola* | EF187937.1 | DQ369570.1 | | DQ369507.1 | | | EF188157.1 |
| *Heterosyphus sicardi* | EF187964.1 | EF188013.1 | | EF188097.1 | | | EF188188.1 |
| *Helictopleurus cribricollis* | EF187922.1 | | | EF188054.1 | | | EF188143.1 |
| *Helictopleurus rudicollis* | EF656675.1 | DQ369580.1 | | FJ817927.1 | | | EF188183.1 |
| *Helictopleurus quadripunctatus* | AY131551.1 | JN619263.1 | | EF656698.1 | | | AY131907.1 |
| *Helictopleurus unifasciatus* | DQ369539.1 | DQ369584.1 | | DQ369521.1 | | | EF188196.1 |
| *Helictopleurus fulgens* | DQ369534.1 | DQ369579.1 | | DQ369516.1 | | | EF188156.1 |
| *Helictopleurus perrieri* | DQ369532.1 | DQ369577.1 | | DQ369514.1 | | | EF188175.1 |
| *Helictopleurus splendidicollis* | DQ369538.1 | DQ369583.1 | | DQ369520.1 | | | EF188191.1 |
| *Helictopleurus fissicollis* | DQ369524.1 | DQ369569.1 | | DQ369506.1 | | | EF188151.1 |
| *Helictopleurus neuter* | EF187951.1 | | | EF188082.1 | | | EF188170.1 |
| *Helictopleurus obscurus* | DQ369531.1 | DQ369576.1 | | DQ369513.1 | | | DQ369458.1 |
| *Helictopleurus multimaculatus* | DQ369528.1 | DQ369573.1 | | DQ369510.1 | | | DQ369455.1 |
| *Helictopleurus fasciolatus* | EF187927.1 | EF187989.1 | | EF188060.1 | | | EF188149.1 |
| *Helictopleurus carbonarius* | EF187918.1 | EF187984.1 | | EF188052.1 | | | EF188140.1 |
| *Helictopleurus marsyas* | DQ369527.1 | DQ369572.1 | | DQ369509.1 | | | EF188163.1 |
| *Helictopleurus nicollei* | DQ369530.1 | DQ369575.1 | | DQ369512.1 | | | EF188172.1 |
| *Helictopleurus corruscus* | DQ369523.1 | DQ369568.1 | | DQ369505.1 | | | EF188141.1 |
| *Helictopleurus sinuaticornis* | DQ369537.1 | DQ369582.1 | | DQ369519.1 | | | EF188189.1 |
| *Helictopleurus giganteus* | DQ369526.1 | DQ369571.1 | | DQ369508.1 | | | EF188161.1 |
| *Helictopleurus steineri* | EF187969.1 | EF188017.1 | | EF656716.1 | | | EF188193.1 |
| *Helictopleurus semivirens* | EF187962.1 | DQ369581.1 | | DQ369518.1 | | | EF188185.1 |
| *Helictopleurus dorbignyi* | EF187925.1 | EF187986.1 | | EF188057.1 | | | EF188146.1 |
| *Helictopleurus minutus* | EF187946.1 | | | EF188078.1 | | | |
| *Helictopleurus viridiflavus* | EF187973.1 | EF188022.1 | | EF188106.1 | | | |

| | | | | | | | |
|---|---|---|---|---|---|---|---|
| *Helictopleurus littoralis* | | | | | | | EU429042.1 |
| *Helictopleurus neoamplicollis* | DQ369529.1 | DQ369574.1 | | DQ369511.1 | | | EF188168.1 |
| *Onthophagus maki* | | | KC294253.1 | | | | KC294242.1 |
| *Onthophagus hirtus* | | | | | | | AY039346.1 |
| *Onthophagus alcyon* | | | | | | | |
| *Hyalonthophagus alcyon* | AY131565.1 | JN619184.1 | | AY131752.1 | | EU162439.1 | AY131920.1 |
| *Onthophagus alcyonides* | EU162533.1 | | | | | | |
| *Hyalonthophagus alcyonides* | JN804706.1 | | JN804777.1 | | | | JN804633.1 |
| *Onthophagus fuliginosus* | EU162543.1 | | | | | KU665398.1 | |
| *Onthophagus auritus* | AY847529.1 | | | | | | |
| *Onthophagus furcaticeps* | AY131576.1 | | | AY131761.1 | | | |
| *Cleptocaccobius convexifrons* | AY131561.1 | JN619210.1 | | AY131748.1 | | | AY131917.1 |
| *Caccobius nigritulus* | AY131559.1 | | | AY131746.1 | | MH020304.1 | AY131915.1 |
| *Caccobius schreberi* | AY131560.1 | | | AY131747.1 | | | AY131916.1 |
| *Onthophagus depressus* | EF187975.1 | EF188032.1 | | EF188116.1 | | | EF188207.1 |
| *Onthophagus hinnulus* | | EF188037.1 | | EF188122.1 | | | EF188214.1 |
| *Onthophagus mije* | AY131581.1 | | | AY131765.1 | | | |
| *Onthophagus ochropygus* | JN804726.1 | | | | | | |
| *Onthophagus fimetarius* | AY131575.1 | | | AY131760.1 | | | AY131925.1 |
| *Onthophagus vulpes* | AY131591.1 | | | AY131775.1 | | | MF944216.1 |
| *Onthophagus pedator* | MF944129.1 | | | | | | MF962927.1 |
| *Onthophagus rorarius* | AY131586.1 | | | AY131770.1 | | | AY131929.1 |
| *Onthophagus orientalis* | MF944127.1 | | | | | | MF944206.1 |
| *Onthophagus maniti* | MF944125.1 | | | | | | MF944204.1 |
| *Euonthophagus carbonarius* | AY131564.1 | JN619193.1 | | AY131751.1 | | | AY131919.1 |
| *Onthophagus vinctus* | | KX512514.1 | KX512603.1 | KX512673.1 | | | |

| | | | | | | | |
|---|---|---|---|---|---|---|---|
| *Onthophagus interstitialis* | JN804698.1 | | JN804770.1 | | JN804575.1 | EU162442.1 | JN804623.1 |
| *Onthophagus binodis* | EU162536.1 | | | | | | |
| *Onthophagus punctatus* | | | | | | | AY039348.1 |
| *Caccobius binodulus* | AY131558.1 | JN619183.1 | | AY131745.1 | | EU162459.1 | AY131914.1 |
| *Onthophagus nigriventris* | EU162555.1 | | | | | EU162475.1 | |
| *Onthophagus sugillatus* | EU162572.1 | | | | | | |
| *Euonthophagus amyntas* | | | | | | | AY039342.1 |
| *Onthophagus borneensis* | MF944122.1 | | | | | | MF944202.1 |
| *Onthophagus furcatus* | | | | | | | AY039357.1 |
| *Onthophagus variegatus* | | EF188040.1 | JN804797.1 | EF188125.1 | | KM441496.1 | EF188217.1 |
| *Onthophagus coenobita* | | | | | | | AY039350.1 |
| *Onthophagus opacicollis* | | | KC294251.1 | | | KM441390.1 | KC294243.1 |
| *Onthophagus fracticornis* | | | KC294254.1 | | | LN832281.1 | KC294245.1 |
| *Onthophagus similis* | AY131590.1 | | KC294256.1 | AY131774.1 | | LN554677.1 | AY131933.1 |
| *Onthophagus lemur* | | | | | | | AY039353.1 |
| *Onthophagus melitaeus* | | | | | | | AY039349.1 |
| *Onthophagus latigena* | | | | | | | AY039356.1 |
| *Onthophagus ruficapillus* | | | KC294255.1 | | | KM445886.1 | KC294246.1 |
| *Onthophagus ovatus* | | | | | | | AY039351.1 |
| *Onthophagus grossepunctatus* | | | | | | KM439793.1 | AY039347.1 |
| *Onthophagus verticicornis* | | | KC294258.1 | | | | KC294249.1 |
| *Onthophagus nutans* | | | | | | | AY039344.1 |
| *Onthophagus stylocerus* | | | | | | LN554680.1 | AY039352.1 |
| *Onthophagus andalusicus* | | | | | | LN832282.1 | |
| *Onthophagus vacca* | | | KC294250.1 | | | KM447997.1 | KC294236.1 |
| *Onthophagus medius* | | | | | | | |

| | | | | | | | |
|---|---|---|---|---|---|---|---|
| *Onthophagus marginalis* | KJ721838.1 | | | KJ721900.1 | | LN554683.1 | |
| *Onthophagus nebulosus* | | | | | | | |
| *Onthophagus merdarius* | | | | | | KU916082.1 | AY039355.1 |
| *Onthophagus nuchicornis* | EU162556.1 | | KC294252.1 | | | | KC294244.1 |
| *Onthophagus glabratus* | AY131577.1 | | | AY131762.1 | | | AY131926.1 |
| *Onthophagus pronus* | AY847525.1 | | | | | EU162451.1 | |
| *Onthophagus granulatus* | EU162545.1 | | | | | | |
| *Onthophagus phanaeides* | MF944130.1 | | | | | | MF944208.1 |
| *Milichus apicalis* | AY131566.1 | JN619196.1 | JN804787.1 | AY131753.1 | | | AY131921.1 |
| *Onthophagus rubicundulus* | AY131587.1 | | | AY131771.1 | | EU162452.1 | AY131930.1 |
| *Onthophagus haagi* | EU162546.1 | | | | | EU162477.1 | |
| *Onthophagus vermiculatus* | EU162575.1 | | | | | | |
| *Onthophagus dunningi* | | | | | | | MG588143.1 |
| *Onthophagus mulgravei* | AY131582.1 | | | AY131766.1 | | | AY131927.1 |
| *Onthophagus turrbal* | AY847531.1 | | | | | | |
| *Onthophagus quadripustulatus* | AY131585.1 | | | AY131769.1 | | | |
| *Onthophagus consentaneus* | AY131573.1 | | | AY131758.1 | | EU162456.1 | |
| *Onthophagus laminatus* | EU162550.1 | | | AY131764.1 | | EU162458.1 | |
| *Onthophagus mjobergi* | EU162553.1 | | | | | EU162468.1 | |
| *Onthophagus sloanei* | EU162565.1 | | | | | EU162449.1 | |
| *Onthophagus ferox* | EU162542.1 | | | | | EU162463.1 | |
| *Onthophagus pentacanthus* | EU162559.1 | | | | | EU162448.1 | |
| *Onthophagus evanidus* | | | | | | KP978430.1 | |
| *Onthophagus ochromerus* | | | | | | | |
| *Onthophagus neostenocerus* | AY847530.1 | | | | | EU162441.1 | |
| *Onthophagus australis* | EU162535.1 | | | | | | |

| | | | | | | | |
|---|---|---|---|---|---|---|---|
| *Onthophagus mammillatus* | AY847526.1 | | | | | | |
| *Onthophagus pugnax* | AY847527.1 | | | | | EU162443.1 | |
| *Onthophagus capella* | EU162537.1 | | | AY131756.1 | | KM450900.1 | |
| *Onthophagus illyricus* | | | | | | | |
| *Onthophagus bivertex* | KJ721867.1 | | | | | EU162476.1 | |
| *Onthophagus taurus* | EU162573.1 | | DQ430885.1 | DQ430937.1 | XM23060308.1 | | KC294248.1 |
| *Onthophagus taurinus* | MF944135.1 | | | | | | MF944213.1 |
| *Onthophagus solivagus* | KJ721822.1 | | | KJ721871.1 | | | JX064146.1 |
| *Onthophagus sinicus* | KJ721837.1 | | | KJ721899.1 | | | |
| *Onthophagus viduus* | KJ721823.1 | | | KJ721872.1 | | | JX064148.1 |
| *Onthophagus obscurior* | AY131584.1 | | | AY131768.1 | | | AY131928.1 |
| *Onthophagus babirussa* | MF944116.1 | | | | | | MF944197.1 |
| *Onthophagus babirussoides* | AY131568.1 | | | AY131754.1 | | EU162440.1 | AY131922.1 |
| *Onthophagus asperulus* | EU162534.1 | | | | | | |
| *Onthophagus fodiens* | KJ721824.1 | | | KJ721873.1 | | | JX064151.1 |
| *Onthophagus lenzi* | JX269021.1 | | | KJ721874.1 | | EU162444.1 | JX064153.1 |
| *Onthophagus clypeatus* | EU162538.1 | | | EF656709.1 | | EU162478.1 | EF656758.1 |
| *Onthophagus xanthomerus* | EU162576.1 | | | | | EU162464.1 | |
| *Onthophagus praecellens* | EU162560.1 | | | | | EU162437.1 | |
| *Onthophagus acuminatus* | EU162531.1 | | | | | EU162474.1 | |
| *Onthophagus stockwelli* | | | | | | | |
| *Onthophagus batesi* | EF656647.1 | | | EF656689.1 | | EU162455.1 | EF656738.1 |
| *Onthophagus incensus* | EU162549.1 | | | | | EU162467.1 | |
| *Onthophagus sharpi* | EU162564.1 | | | | | | |
| *Onthophagus championi* | EF656651.1 | | | EF656693.1 | | | EF656742.1 |
| *Onthophagus bidentatus* | AY131569.1 | | | AY131755.1 | | EU162457.1 | AY131923.1 |

| | | | | | | | |
|---|---|---|---|---|---|---|---|
| *Onthophagus marginicollis* | EU162552.1 | | | | | EU162453.1 | |
| *Onthophagus haematopus* | EU162547.1 | | | EF656711.1 | | MG059101.1 | EF656760.1 |
| *Onthophagus orpheus* | EU162557.1 | | | | | EU162454.1 | |
| *Onthophagus hecate* | EU162548.1 | | DQ430887.1 | DQ430939.1 | | EU162445.1 | |
| *Onthophagus coscineus* | EU162539.1 | | | | | EU162447.1 | |
| *Onthophagus crinitus* | EU162541.1 | AY821535.1 | | AY131759.1 | | EU162462.1 | AY131924.1 |
| *Onthophagus pennsylvanicus* | EU162558.1 | | | | | | |

**Table S2 Generic Monophyly Dung Beetles.**

Monophyly for all genera represented in the phylogeny, assessed for the original data set and each of the trees made from the bootstrapped data respectively. The last three columns sum how often a genus was monophyletic ("Yes") or non-monophyletic ("No") across all trees. Monot.=Monotypic.

| Genus | Full | BS0 | BS1 | BS2 | BS3 | BS4 | BS5 | BS6 | BS7 | BS8 | BS9 | Yes | No |
|---|---|---|---|---|---|---|---|---|---|---|---|---|---|
| *Byrrhidium* | Yes | Yes | Yes | Yes | Yes | Yes | Yes | Yes | Yes | Yes | Yes | 11 | 0 |
| *Catharsius* | Yes | Yes | Yes | Yes | Yes | Yes | Yes | Yes | Yes | Yes | Yes | 11 | 0 |
| *Dicranocara* | Yes | Yes | Yes | Yes | Yes | Yes | Yes | Yes | Yes | Yes | Yes | 11 | 0 |
| *Eucranium* | Yes | Yes | Yes | Yes | Yes | Yes | Yes | Yes | Yes | Yes | Yes | 11 | 0 |
| *Eurysternus* | Yes | Yes | Yes | Yes | Yes | Yes | Yes | Yes | Yes | Yes | Yes | 11 | 0 |
| *Gymnopleurus* | Yes | Yes | Yes | Yes | Yes | Yes | Yes | Yes | Yes | Yes | Yes | 11 | 0 |
| *Heliocopris* | Yes | Yes | Yes | Yes | Yes | Yes | Yes | Yes | Yes | Yes | Yes | 11 | 0 |
| *Kheper* | Yes | Yes | Yes | Yes | Yes | Yes | Yes | Yes | Yes | Yes | Yes | 11 | 0 |
| *Macroderes* | Yes | Yes | Yes | Yes | Yes | Yes | Yes | Yes | Yes | Yes | Yes | 11 | 0 |
| *Metacatharsius* | Yes | Yes | Yes | Yes | Yes | Yes | Yes | Yes | Yes | Yes | Yes | 11 | 0 |
| *Pachysoma* | Yes | Yes | Yes | Yes | Yes | Yes | Yes | Yes | Yes | Yes | Yes | 11 | 0 |
| *Paragymnopleurus* | Yes | Yes | Yes | Yes | Yes | Yes | Yes | Yes | Yes | Yes | Yes | 11 | 0 |
| *Sarophorus* | Yes | Yes | Yes | Yes | Yes | Yes | Yes | Yes | Yes | Yes | Yes | 11 | 0 |
| *Xinidium* | Yes | Yes | Yes | Yes | Yes | Yes | Yes | Yes | Yes | Yes | Yes | 11 | 0 |
| *Ateuchus* | Yes | Yes | Yes | Yes | No | Yes | Yes | Yes | Yes | Yes | Yes | 10 | 1 |
| *Cheironitis* | Yes | Yes | Yes | Yes | No | Yes | Yes | Yes | Yes | Yes | Yes | 10 | 1 |
| *Coptodactyla* | Yes | Yes | Yes | Yes | Yes | Yes | No | Yes | Yes | Yes | Yes | 10 | 1 |
| *Epactoides* | Yes | Yes | Yes | Yes | Yes | Yes | No | Yes | Yes | Yes | Yes | 10 | 1 |
| *Epirinus* | Yes | Yes | Yes | Yes | Yes | No | Yes | Yes | Yes | Yes | Yes | 10 | 1 |
| *Sauvagesinella* | Yes | Yes | Yes | Yes | Yes | Yes | Yes | Yes | Yes | No | Yes | 10 | 1 |

| | | | | | | | | | | | | | |
|---|---|---|---|---|---|---|---|---|---|---|---|---|---|
| *Anachalcos* | Yes | Yes | No | Yes | Yes | Yes | Yes | Yes | Yes | No | Yes | 9 | 2 |
| *Demarziella* | Yes | Yes | Yes | Yes | No | Yes | No | Yes | Yes | Yes | Yes | 9 | 2 |
| *Lepanus* | Yes | No | Yes | Yes | Yes | No | Yes | Yes | Yes | Yes | Yes | 9 | 2 |
| *Saphobius* | Yes | Yes | Yes | No | Yes | Yes | Yes | No | Yes | Yes | Yes | 9 | 2 |
| *Cephalodesmius* | Yes | No | Yes | Yes | No | No | Yes | Yes | Yes | Yes | Yes | 8 | 3 |
| *Pseudoarachnodes* | Yes | Yes | Yes | Yes | No | Yes | Yes | Yes | No | No | Yes | 8 | 3 |
| *Scybalophagus* | Yes | Yes | No | Yes | Yes | No | Yes | Yes | No | Yes | Yes | 8 | 3 |
| *Temnoplectron* | Yes | No | Yes | Yes | No | Yes | No | Yes | Yes | Yes | Yes | 8 | 3 |
| *Uroxys* | No | Yes | No | Yes | Yes | Yes | Yes | No | Yes | Yes | Yes | 8 | 3 |
| *Deltochilum* | Yes | No | Yes | Yes | Yes | Yes | Yes | Yes | No | No | No | 7 | 4 |
| *Canthonosoma* | Yes | Yes | Yes | Yes | No | No | No | No | No | No | Yes | 5 | 6 |
| *Euoniticellus* | Yes | No | Yes | Yes | Yes | Yes | No | No | No | No | No | 5 | 6 |
| *Dichotomius* | Yes | Yes | No | No | No | No | Yes | No | No | Yes | No | 4 | 7 |
| *Bubas* | Yes | No | No | Yes | No | No | Yes | No | No | No | No | 3 | 8 |
| *Diorygopyx* | No | Yes | Yes | No | No | No | No | No | No | Yes | No | 3 | 8 |
| *Phalops* | No | No | Yes | No | Yes | Yes | No | No | No | No | No | 3 | 8 |
| *Canthidium* | No | Yes | No | Yes | No | No | No | No | No | No | No | 2 | 9 |
| *Drepanopodus* | No | No | No | No | No | No | No | Yes | No | No | No | 1 | 10 |
| *Glyphoderus* | No | No | No | No | No | No | Yes | No | No | No | No | 1 | 10 |
| *Aliuscanthoniola* | Monot. | Monot. | Monot. | Monot. | Monot. | Monot. | Monot. | Monot. | Monot. | Monot. | Monot. | 0 | 0 |
| *Allogymnopleurus* | Monot. | Monot. | Monot. | Monot. | Monot. | Monot. | Monot. | Monot. | Monot. | Monot. | Monot. | 0 | 0 |
| *Amphistomus* | Monot. | Monot. | Monot. | Monot. | Monot. | Monot. | Monot. | Monot. | Monot. | Monot. | Monot. | 0 | 0 |
| *Anonthobium* | Monot. | Monot. | Monot. | Monot. | Monot. | Monot. | Monot. | Monot. | Monot. | Monot. | Monot. | 0 | 0 |
| *Bdelyropsis* | Monot. | Monot. | Monot. | Monot. | Monot. | Monot. | Monot. | Monot. | Monot. | Monot. | Monot. | 0 | 0 |
| *Bohepilissus* | Monot. | Monot. | Monot. | Monot. | Monot. | Monot. | Monot. | Monot. | Monot. | Monot. | Monot. | 0 | 0 |

| | | | | | | | | | | | | | |
|---|---|---|---|---|---|---|---|---|---|---|---|---|---|
| *Boletoscapter* | Monot. | Monot. | Monot. | Monot. | Monot. | Monot. | Monot. | Monot. | Monot. | Monot. | Monot. | 0 | 0 |
| *Cambefortantus* | Monot. | Monot. | Monot. | Monot. | Monot. | Monot. | Monot. | Monot. | Monot. | Monot. | Monot. | 0 | 0 |
| *Canthodimorpha* | Monot. | Monot. | Monot. | Monot. | Monot. | Monot. | Monot. | Monot. | Monot. | Monot. | Monot. | 0 | 0 |
| *Cleptocaccobius* | Monot. | Monot. | Monot. | Monot. | Monot. | Monot. | Monot. | Monot. | Monot. | Monot. | Monot. | 0 | 0 |
| *Coproecus* | Monot. | Monot. | Monot. | Monot. | Monot. | Monot. | Monot. | Monot. | Monot. | Monot. | Monot. | 0 | 0 |
| *Coptorhina* | Monot. | Monot. | Monot. | Monot. | Monot. | Monot. | Monot. | Monot. | Monot. | Monot. | Monot. | 0 | 0 |
| *Cyptochirus* | Monot. | Monot. | Monot. | Monot. | Monot. | Monot. | Monot. | Monot. | Monot. | Monot. | Monot. | 0 | 0 |
| *Delopleurus* | Monot. | Monot. | Monot. | Monot. | Monot. | Monot. | Monot. | Monot. | Monot. | Monot. | Monot. | 0 | 0 |
| *Dendropaemon* | Monot. | Monot. | Monot. | Monot. | Monot. | Monot. | Monot. | Monot. | Monot. | Monot. | Monot. | 0 | 0 |
| *Diabroctis* | Monot. | Monot. | Monot. | Monot. | Monot. | Monot. | Monot. | Monot. | Monot. | Monot. | Monot. | 0 | 0 |
| *Digitonthophagus* | Monot. | Monot. | Monot. | Monot. | Monot. | Monot. | Monot. | Monot. | Monot. | Monot. | Monot. | 0 | 0 |
| *Dwesasilvasedis* | Monot. | Monot. | Monot. | Monot. | Monot. | Monot. | Monot. | Monot. | Monot. | Monot. | Monot. | 0 | 0 |
| *Endroedyolus* | Monot. | Monot. | Monot. | Monot. | Monot. | Monot. | Monot. | Monot. | Monot. | Monot. | Monot. | 0 | 0 |
| *Ennearabdus* | Monot. | Monot. | Monot. | Monot. | Monot. | Monot. | Monot. | Monot. | Monot. | Monot. | Monot. | 0 | 0 |
| *Eodrepanus* | Monot. | Monot. | Monot. | Monot. | Monot. | Monot. | Monot. | Monot. | Monot. | Monot. | Monot. | 0 | 0 |
| *Eudinopus* | Monot. | Monot. | Monot. | Monot. | Monot. | Monot. | Monot. | Monot. | Monot. | Monot. | Monot. | 0 | 0 |
| *Frankenbergerius* | Monot. | Monot. | Monot. | Monot. | Monot. | Monot. | Monot. | Monot. | Monot. | Monot. | Monot. | 0 | 0 |
| *Garreta* | Monot. | Monot. | Monot. | Monot. | Monot. | Monot. | Monot. | Monot. | Monot. | Monot. | Monot. | 0 | 0 |
| *Gyronotus* | Monot. | Monot. | Monot. | Monot. | Monot. | Monot. | Monot. | Monot. | Monot. | Monot. | Monot. | 0 | 0 |
| *Hammondantus* | Monot. | Monot. | Monot. | Monot. | Monot. | Monot. | Monot. | Monot. | Monot. | Monot. | Monot. | 0 | 0 |
| *Hansreia* | Monot. | Monot. | Monot. | Monot. | Monot. | Monot. | Monot. | Monot. | Monot. | Monot. | Monot. | 0 | 0 |
| *Heterosyphus* | Monot. | Monot. | Monot. | Monot. | Monot. | Monot. | Monot. | Monot. | Monot. | Monot. | Monot. | 0 | 0 |
| *Ignambia* | Monot. | Monot. | Monot. | Monot. | Monot. | Monot. | Monot. | Monot. | Monot. | Monot. | Monot. | 0 | 0 |
| *Janssensantus* | Monot. | Monot. | Monot. | Monot. | Monot. | Monot. | Monot. | Monot. | Monot. | Monot. | Monot. | 0 | 0 |
| *Leotrichillum* | Monot. | Monot. | Monot. | Monot. | Monot. | Monot. | Monot. | Monot. | Monot. | Monot. | Monot. | 0 | 0 |

| | | | | | | | | | | | | | |
|---|---|---|---|---|---|---|---|---|---|---|---|---|---|
| *Litocopris* | Monot. | Monot. | Monot. | Monot. | Monot. | Monot. | Monot. | Monot. | Monot. | Monot. | Monot. | 0 | 0 |
| *Malagoniella* | Monot. | Monot. | Monot. | Monot. | Monot. | Monot. | Monot. | Monot. | Monot. | Monot. | Monot. | 0 | 0 |
| *Megathopa* | Monot. | Monot. | Monot. | Monot. | Monot. | Monot. | Monot. | Monot. | Monot. | Monot. | Monot. | 0 | 0 |
| *Megathoposoma* | Monot. | Monot. | Monot. | Monot. | Monot. | Monot. | Monot. | Monot. | Monot. | Monot. | Monot. | 0 | 0 |
| *Milichus* | Monot. | Monot. | Monot. | Monot. | Monot. | Monot. | Monot. | Monot. | Monot. | Monot. | Monot. | 0 | 0 |
| *Monoplistes* | Monot. | Monot. | Monot. | Monot. | Monot. | Monot. | Monot. | Monot. | Monot. | Monot. | Monot. | 0 | 0 |
| *Monteithocanthon* | Monot. | Monot. | Monot. | Monot. | Monot. | Monot. | Monot. | Monot. | Monot. | Monot. | Monot. | 0 | 0 |
| *Namakwanus* | Monot. | Monot. | Monot. | Monot. | Monot. | Monot. | Monot. | Monot. | Monot. | Monot. | Monot. | 0 | 0 |
| *Neateuchus* | Monot. | Monot. | Monot. | Monot. | Monot. | Monot. | Monot. | Monot. | Monot. | Monot. | Monot. | 0 | 0 |
| *Neosisyphus* | Monot. | Monot. | Monot. | Monot. | Monot. | Monot. | Monot. | Monot. | Monot. | Monot. | Monot. | 0 | 0 |
| *Ochicanthon* | Monot. | Monot. | Monot. | Monot. | Monot. | Monot. | Monot. | Monot. | Monot. | Monot. | Monot. | 0 | 0 |
| *Odontoloma* | Monot. | Monot. | Monot. | Monot. | Monot. | Monot. | Monot. | Monot. | Monot. | Monot. | Monot. | 0 | 0 |
| *Onthobium* | Monot. | Monot. | Monot. | Monot. | Monot. | Monot. | Monot. | Monot. | Monot. | Monot. | Monot. | 0 | 0 |
| *Outenikwanus* | Monot. | Monot. | Monot. | Monot. | Monot. | Monot. | Monot. | Monot. | Monot. | Monot. | Monot. | 0 | 0 |
| *Pachylomera* | Monot. | Monot. | Monot. | Monot. | Monot. | Monot. | Monot. | Monot. | Monot. | Monot. | Monot. | 0 | 0 |
| *Paracopris* | Monot. | Monot. | Monot. | Monot. | Monot. | Monot. | Monot. | Monot. | Monot. | Monot. | Monot. | 0 | 0 |
| *Paraphytus* | Monot. | Monot. | Monot. | Monot. | Monot. | Monot. | Monot. | Monot. | Monot. | Monot. | Monot. | 0 | 0 |
| *Parascatonomus* | Monot. | Monot. | Monot. | Monot. | Monot. | Monot. | Monot. | Monot. | Monot. | Monot. | Monot. | 0 | 0 |
| *Paronthobium* | Monot. | Monot. | Monot. | Monot. | Monot. | Monot. | Monot. | Monot. | Monot. | Monot. | Monot. | 0 | 0 |
| *Peckolus* | Monot. | Monot. | Monot. | Monot. | Monot. | Monot. | Monot. | Monot. | Monot. | Monot. | Monot. | 0 | 0 |
| *Pseudignambia* | Monot. | Monot. | Monot. | Monot. | Monot. | Monot. | Monot. | Monot. | Monot. | Monot. | Monot. | 0 | 0 |
| *Pseudonthobium* | Monot. | Monot. | Monot. | Monot. | Monot. | Monot. | Monot. | Monot. | Monot. | Monot. | Monot. | 0 | 0 |
| *Pycnopanelus* | Monot. | Monot. | Monot. | Monot. | Monot. | Monot. | Monot. | Monot. | Monot. | Monot. | Monot. | 0 | 0 |
| *Scybalocanthon* | Monot. | Monot. | Monot. | Monot. | Monot. | Monot. | Monot. | Monot. | Monot. | Monot. | Monot. | 0 | 0 |
| *Serrophorus* | Monot. | Monot. | Monot. | Monot. | Monot. | Monot. | Monot. | Monot. | Monot. | Monot. | Monot. | 0 | 0 |

| | | | | | | | | | | | | |
|---|---|---|---|---|---|---|---|---|---|---|---|---|
| *Silvaphilus* | Monot. | Monot. | Monot. | Monot. | Monot. | Monot. | Monot. | Monot. | Monot. | Monot. | Monot. | 0 | 0 |
| *Sulcophanaeus* | Monot. | Monot. | Monot. | Monot. | Monot. | Monot. | Monot. | Monot. | Monot. | Monot. | Monot. | 0 | 0 |
| *Tanzanolus* | Monot. | Monot. | Monot. | Monot. | Monot. | Monot. | Monot. | Monot. | Monot. | Monot. | Monot. | 0 | 0 |
| *Tragiscus* | Monot. | Monot. | Monot. | Monot. | Monot. | Monot. | Monot. | Monot. | Monot. | Monot. | Monot. | 0 | 0 |
| *Trichillidium* | Monot. | Monot. | Monot. | Monot. | Monot. | Monot. | Monot. | Monot. | Monot. | Monot. | Monot. | 0 | 0 |
| *Anomiopsoides* | No | No | No | No | No | No | No | No | No | No | No | 0 | 11 |
| *Apotolamprus* | No | No | No | No | No | No | No | No | No | No | No | 0 | 11 |
| *Arachnodes* | No | No | No | No | No | No | No | No | No | No | No | 0 | 11 |
| *Caccobius* | No | No | No | No | No | No | No | No | No | No | No | 0 | 11 |
| *Canthon* | No | No | No | No | No | No | No | No | No | No | No | 0 | 11 |
| *Copris* | No | No | No | No | No | No | No | No | No | No | No | 0 | 11 |
| *Coprophanaeus* | No | No | No | No | No | No | No | No | No | No | No | 0 | 11 |
| *Drepanocerus* | No | No | No | No | No | No | No | No | No | No | No | 0 | 11 |
| *Euonthophagus* | No | No | No | No | No | No | No | No | No | No | No | 0 | 11 |
| *Helictopleurus* | No | No | No | No | No | No | No | No | No | No | No | 0 | 11 |
| *Hyalonthophagus* | No | No | No | No | No | No | No | No | No | No | No | 0 | 11 |
| *Liatongus* | No | No | No | No | No | No | No | No | No | No | No | 0 | 11 |
| *Nanos* | No | No | No | No | No | No | No | No | No | No | No | 0 | 11 |
| *Oniticellus* | No | No | No | No | No | No | No | No | No | No | No | 0 | 11 |
| *Onitis* | No | No | No | No | No | No | No | No | No | No | No | 0 | 11 |
| *Onthophagus* | No | No | No | No | No | No | No | No | No | No | No | 0 | 11 |
| *Oxysternon* | No | No | No | No | No | No | No | No | No | No | No | 0 | 11 |
| *Phanaeus* | No | No | No | No | No | No | No | No | No | No | No | 0 | 11 |
| *Proagoderus* | No | No | No | No | No | No | No | No | No | No | No | 0 | 11 |
| *Scarabaeus* | No | No | No | No | No | No | No | No | No | No | No | 0 | 11 |

| | | | | | | | | | | | | | |
|---|---|---|---|---|---|---|---|---|---|---|---|---|---|
| *Sisyphus* | No | No | No | No | No | No | No | No | No | No | No | 0 | 11 |
| *Tiniocellus* | No | No | No | No | No | No | No | No | No | No | No | 0 | 11 |

**Table S3 Tribal Monophyly Dung Beetles.**

Monophyly for all tribes represented in the phylogeny, assessed for the original data set and each of the trees made from the bootstrapped data respectively.

| Tribe | Full | BS0 | BS1 | BS2 | BS3 | BS4 | BS5 | BS6 | BS7 | BS8 | BS9 |
|---|---|---|---|---|---|---|---|---|---|---|---|
| Phanaeini | No | No | No | No | No | No | No | No | No | No | No |
| Dichotomiini | No | No | No | No | No | No | No | No | No | No | No |
| Eucraniini | Yes | Yes | No | Yes | Yes | Yes | Yes | Yes | Yes | Yes | Yes |
| Deltochilini | No | No | No | No | No | No | No | No | No | No | No |
| Canthonini | No | No | No | No | No | No | No | No | No | No | No |
| Coprini | No | No | No | No | No | No | No | No | No | No | No |
| Ateuchini | Yes | Yes | No | Yes | Yes | Yes | No | No | Yes | No | No |
| Scarabaeini | No | No | No | No | No | No | No | No | No | No | No |
| Eurysternini | Yes | Yes | Yes | Yes | Yes | Yes | Yes | Yes | Yes | Yes | Yes |
| Gymnopleurini | Yes | Yes | Yes | Yes | Yes | Yes | Yes | Yes | Yes | Yes | Yes |
| Onthophagini | No | No | No | No | No | No | No | No | No | No | No |
| Oniticellini | No | No | No | No | No | No | No | No | No | No | No |
| Onitini | No | Yes | No | No | No | No | Yes | No | No | No | Yes |
| Sisyphini | Yes | Yes | Yes | Yes | Yes | Yes | Yes | Yes | Yes | Yes | Yes |

**Table S4 Manual Dispersal Rate Multiplier Matrices.**

Manual dispersal rates for dispersal between all areas, for each defined epoch (duration see header row). The colors represent large sea barriers (dark blue), short sea barriers (light blue), land barriers (yellow), direct adjacency (green), and maximal barrier strength (light gray). Af=Africa, As=Asia, Eu=Europe, Na=North America, Oc= Oceania, Sa=South America, Ma=Madagascar.

| **20-0 ma** | | | | | | | |
|---|---|---|---|---|---|---|---|
| | Af | As | Eu | Na | Oc | Sa | Ma |
| Af | 1 | 0.85 | 0.85 | 0.25 | 0.25 | 0.25 | 0.75 |
| As | 0.85 | 1 | 1 | 0.85 | 0.85 | 0.25 | 0.25 |
| Eu | 0.85 | 1 | 1 | 0.75 | 0.05 | 0.25 | 0.25 |
| Na | 0.25 | 0.85 | 0.75 | 1 | 0.05 | 0.85 | 0.25 |
| Oc | 0.25 | 0.85 | 0.05 | 0.05 | 1 | 0.25 | 0.25 |
| Sa | 0.25 | 0.25 | 0.25 | 0.85 | 0.25 | 1 | 0.25 |
| Ma | 0.75 | 0.25 | 0.25 | 0.25 | 0.25 | 0.25 | 1 |

| **50-20 ma** | | | | | | | |
|---|---|---|---|---|---|---|---|
| | Af | As | Eu | Na | Oc | Sa | Ma |
| Af | 1 | 0.75 | 0.85 | 0.25 | 0.25 | 0.25 | 0.75 |
| As | 0.75 | 1 | 1 | 0.85 | 0.25 | 0.25 | 0.25 |
| Eu | 0.85 | 1 | 1 | 0.75 | 0.05 | 0.25 | 0.25 |
| Na | 0.25 | 0.85 | 0.75 | 1 | 0.05 | 0.75 | 0.25 |
| Oc | 0.25 | 0.25 | 0.05 | 0.05 | 1 | 0.25 | 0.25 |
| Sa | 0.25 | 0.25 | 0.25 | 0.75 | 0.25 | 1 | 0.25 |
| Ma | 0.75 | 0.25 | 0.25 | 0.25 | 0.25 | 0.25 | 1 |

| **110-50 ma** | | | | | | | |
|---|---|---|---|---|---|---|---|
| | Af | As | Eu | Na | Oc | Sa | Ma |
| Af | 1 | 0.35 | 0.85 | 0.35 | 0.25 | 0.75 | 0.75 |
| As | 0.35 | 1 | 1 | 1 | 0.25 | 0.25 | 0.25 |
| Eu | 0.85 | 1 | 1 | 1 | 0.25 | 0.25 | 0.25 |
| Na | 0.35 | 1 | 1 | 1 | 0.05 | 0.75 | 0.25 |
| Oc | 0.25 | 0.25 | 0.25 | 0.05 | 1 | 0.35 | 0.25 |
| Sa | 0.75 | 0.25 | 0.25 | 0.75 | 0.35 | 1 | 0.25 |
| Ma | 0.75 | 0.25 | 0.25 | 0.25 | 0.25 | 0.25 | 1 |

| **150-110 ma** | | | | | | | |
|---|---|---|---|---|---|---|---|
| | Af | As | Eu | Na | Oc | Sa | Ma |
| Af | 1 | 0.35 | 0.85 | 0.75 | 0.5 | 1 | 0.75 |
| As | 0.35 | 1 | 1 | 1 | 0.25 | 0.35 | 0.25 |
| Eu | 0.85 | 1 | 1 | 1 | 0.25 | 0.35 | 0.25 |
| Na | 0.75 | 1 | 1 | 1 | 0.05 | 0.85 | 0.05 |
| Oc | 0.5 | 0.25 | 0.25 | 0.05 | 1 | 0.5 | 1 |
| Sa | 1 | 0.35 | 0.35 | 0.85 | 0.5 | 1 | 0.5 |
| Ma | 0.75 | 0.25 | 0.25 | 0.05 | 1 | 0.5 | 1 |

| **200-150 ma** | | | | | | | |
|---|---|---|---|---|---|---|---|
| | Af | As | Eu | Na | Oc | Sa | Ma |
| Af | 1 | 0.5 | 1 | 1 | 0.5 | 1 | 1 |
| As | 0.5 | 1 | 1 | 1 | 0.25 | 0.5 | 0.25 |
| Eu | 1 | 1 | 1 | 1 | 0.25 | 0.5 | 0.5 |
| Na | 1 | 1 | 1 | 1 | 0.25 | 1 | 0.5 |
| Oc | 0.5 | 0.25 | 0.25 | 0.25 | 1 | 0.5 | 1 |
| Sa | 1 | 0.5 | 0.5 | 1 | 0.5 | 1 | 0.5 |
| Ma | 1 | 0.25 | 0.5 | 0.5 | 1 | 0.5 | 1 |

**Table S5 Averaged Manual Dispersal Rate Multiplier Matrices.**

Manual dispersal rates for dispersal between all areas, averaged over all epochs (for 4 or 3 epochs, see header row). Af=Africa, As=Asia, Eu=Europe, Na=North America, Oc= Oceania, Sa=South America, Ma=Madagascar.

| **4-Epoch averaged** | | | | | | | |
|---|---|---|---|---|---|---|---|
| | Af | As | Eu | Na | Oc | Sa | Ma |
| Af | 1.00 | 0.66 | 0.85 | 0.33 | 0.28 | 0.45 | 0.75 |
| As | 0.66 | 1.00 | 1.00 | 0.90 | 0.50 | 0.26 | 0.25 |
| Eu | 0.85 | 1.00 | 1.00 | 0.83 | 0.12 | 0.26 | 0.25 |
| Na | 0.33 | 0.90 | 0.83 | 1.00 | 0.05 | 0.80 | 0.23 |
| Oc | 0.28 | 0.50 | 0.12 | 0.05 | 1.00 | 0.30 | 0.33 |
| Sa | 0.45 | 0.26 | 0.26 | 0.80 | 0.30 | 1.00 | 0.28 |
| Ma | 0.75 | 0.25 | 0.25 | 0.23 | 0.33 | 0.28 | 1.00 |

| **3-Epoch averaged** | | | | | | | |
|---|---|---|---|---|---|---|---|
| | Af | As | Eu | Na | Oc | Sa | Ma |
| Af | 1.00 | 0.69 | 0.85 | 0.28 | 0.25 | 0.38 | 0.75 |
| As | 0.69 | 1.00 | 1.00 | 0.89 | 0.53 | 0.25 | 0.25 |
| Eu | 0.85 | 1.00 | 1.00 | 0.81 | 0.10 | 0.25 | 0.25 |
| Na | 0.28 | 0.89 | 0.81 | 1.00 | 0.05 | 0.80 | 0.25 |
| Oc | 0.25 | 0.53 | 0.10 | 0.05 | 1.00 | 0.28 | 0.25 |
| Sa | 0.38 | 0.25 | 0.25 | 0.80 | 0.28 | 1.00 | 0.25 |
| Ma | 0.75 | 0.25 | 0.25 | 0.25 | 0.25 | 0.25 | 1.00 |

**Table S6 BioGeoBEARS Root Range Probabilities.**

Probabilities of the ten ancestral ranges with the on average highest relative probability at the root node, for each tree and bootstrap tree. The most probable area for each tree is highlighted in green. Af=Africa, Oc=Oceania, Sa=South America, Ma=Madagascar, As=Asia, Na=North America.

| Tree | AfOcSa | AfSaMa | AfOc | AfSa | AfOcMa | Af | AfAsOc | AfNaSa | AfNaOc | AfMa |
|---|---|---|---|---|---|---|---|---|---|---|
| Young | 0.9184 | 0.0001 | 0.0129 | 0.0039 | 0.0006 | 0.0001 | 0.0002 | 0.0000 | 0.0634 | 0.0000 |
| Old | 0.9170 | 0.0001 | 0.0122 | 0.0030 | 0.0008 | 0.0001 | 0.0002 | 0.0000 | 0.0663 | 0.0000 |
| Young BS0 | 0.2215 | 0.0000 | 0.6561 | 0.0007 | 0.1016 | 0.0066 | 0.0024 | 0.0000 | 0.0099 | 0.0003 |
| Young BS1 | 0.0577 | 0.6080 | 0.0008 | 0.2930 | 0.0013 | 0.0138 | 0.0000 | 0.0077 | 0.0027 | 0.0073 |
| Young BS2 | 0.8633 | 0.0002 | 0.0913 | 0.0146 | 0.0081 | 0.0127 | 0.0003 | 0.0018 | 0.0019 | 0.0002 |
| Young BS3 | 0.8369 | 0.0010 | 0.0347 | 0.0888 | 0.0002 | 0.0301 | 0.0003 | 0.0021 | 0.0002 | 0.0003 |
| Young BS4 | 0.9573 | 0.0010 | 0.0195 | 0.0121 | 0.0003 | 0.0024 | 0.0001 | 0.0002 | 0.0065 | 0.0000 |
| Young BS5 | 0.0300 | 0.5061 | 0.0128 | 0.0189 | 0.2487 | 0.0312 | 0.0210 | 0.0001 | 0.0003 | 0.0629 |
| Young BS6 | 0.9183 | 0.0001 | 0.0132 | 0.0015 | 0.0021 | 0.0001 | 0.0041 | 0.0000 | 0.0602 | 0.0000 |
| Young BS7 | 0.3352 | 0.3389 | 0.0082 | 0.1850 | 0.0192 | 0.0256 | 0.0001 | 0.0269 | 0.0267 | 0.0065 |
| Young BS8 | 0.0020 | 0.0000 | 0.2655 | 0.0003 | 0.3904 | 0.0534 | 0.2495 | 0.0000 | 0.0067 | 0.0054 |
| Young BS9 | 0.2381 | 0.4228 | 0.0030 | 0.1638 | 0.0000 | 0.0111 | 0.0000 | 0.1298 | 0.0002 | 0.0043 |
| Old BS0 | 0.1342 | 0.0000 | 0.7518 | 0.0017 | 0.0610 | 0.0329 | 0.0037 | 0.0000 | 0.0112 | 0.0010 |
| Old BS1 | 0.0979 | 0.5331 | 0.0040 | 0.2713 | 0.0033 | 0.0438 | 0.0000 | 0.0086 | 0.0044 | 0.0198 |
| Old BS2 | 0.6501 | 0.0009 | 0.1738 | 0.0430 | 0.0132 | 0.0873 | 0.0012 | 0.0055 | 0.0049 | 0.0013 |
| Old BS3 | 0.7257 | 0.0020 | 0.0467 | 0.1317 | 0.0003 | 0.0772 | 0.0004 | 0.0040 | 0.0003 | 0.0009 |
| Old BS4 | 0.8966 | 0.0013 | 0.0509 | 0.0226 | 0.0008 | 0.0127 | 0.0002 | 0.0003 | 0.0132 | 0.0002 |
| Old BS5 | 0.0151 | 0.4344 | 0.0156 | 0.0172 | 0.2953 | 0.0394 | 0.0218 | 0.0000 | 0.0002 | 0.0829 |
| Old BS6 | 0.9002 | 0.0002 | 0.0279 | 0.0027 | 0.0049 | 0.0003 | 0.0085 | 0.0000 | 0.0540 | 0.0001 |
| Old BS7 | 0.3453 | 0.2859 | 0.0148 | 0.1946 | 0.0114 | 0.0532 | 0.0003 | 0.0218 | 0.0241 | 0.0108 |
| Old BS8 | 0.0020 | 0.0001 | 0.2837 | 0.0010 | 0.3020 | 0.1604 | 0.1803 | 0.0000 | 0.0048 | 0.0160 |
| Old BS9 | 0.2318 | 0.3596 | 0.0041 | 0.1849 | 0.0000 | 0.0194 | 0.0000 | 0.1634 | 0.0001 | 0.0054 |

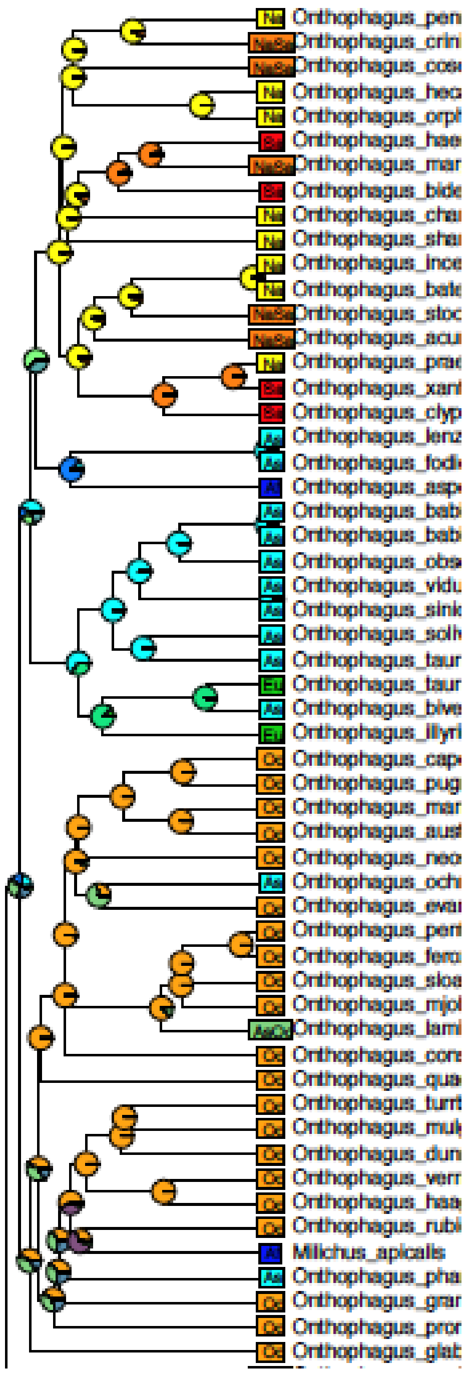


**Figure S1: BioGeoBEARS Ancestral Range Estimation Young Tree.**

Relative probabilities of estimated ancestral ranges at each node indicated by pie charts. Model: DEC+w. Af=Africa, Oc=Oceania, Sa=South America, Ma=Madagascar, As=Asia, Na=North America.

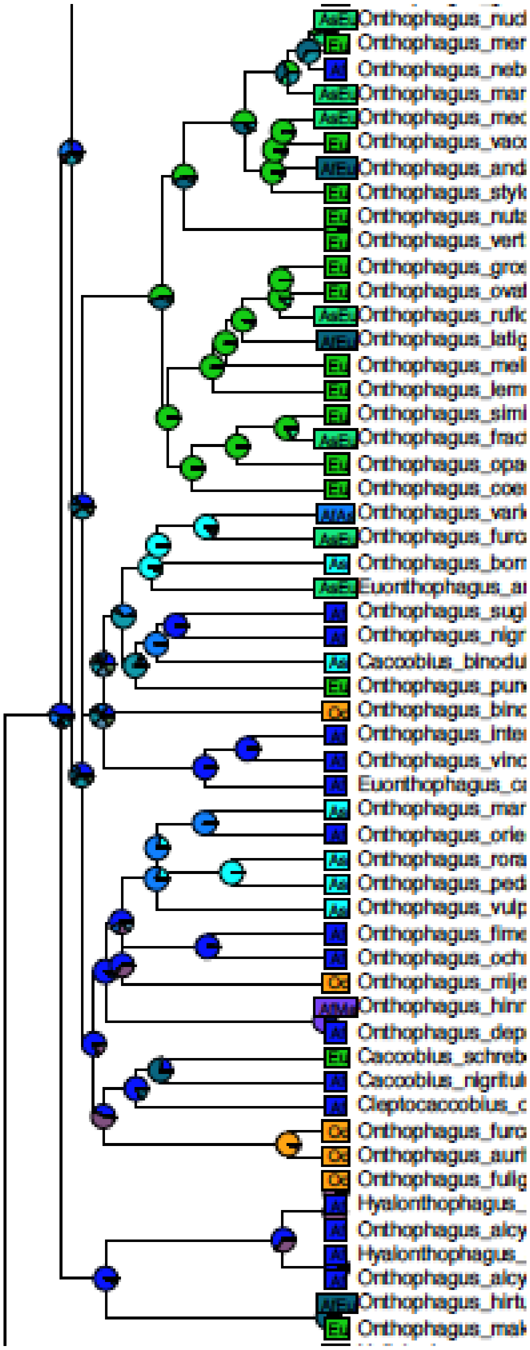


Figure S1 Continued

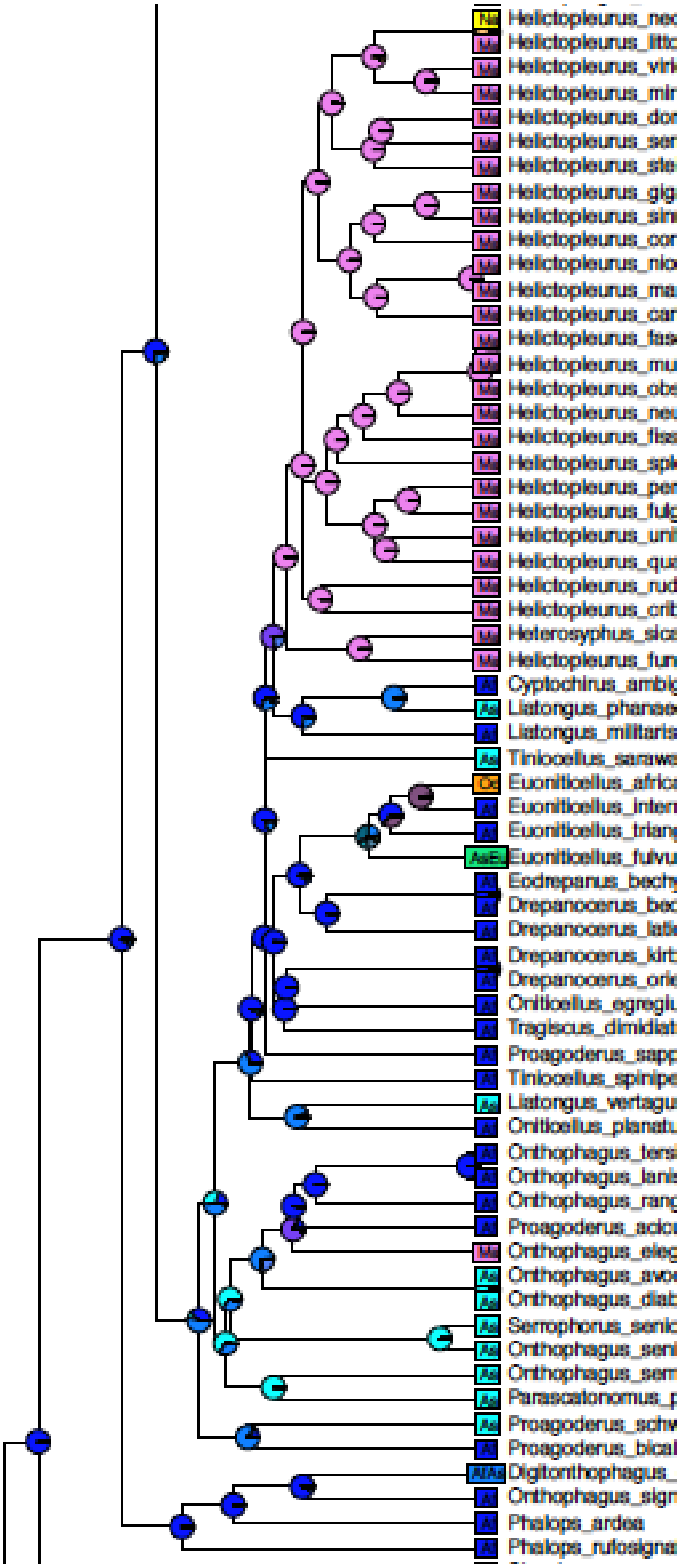


Figure S1 Continued

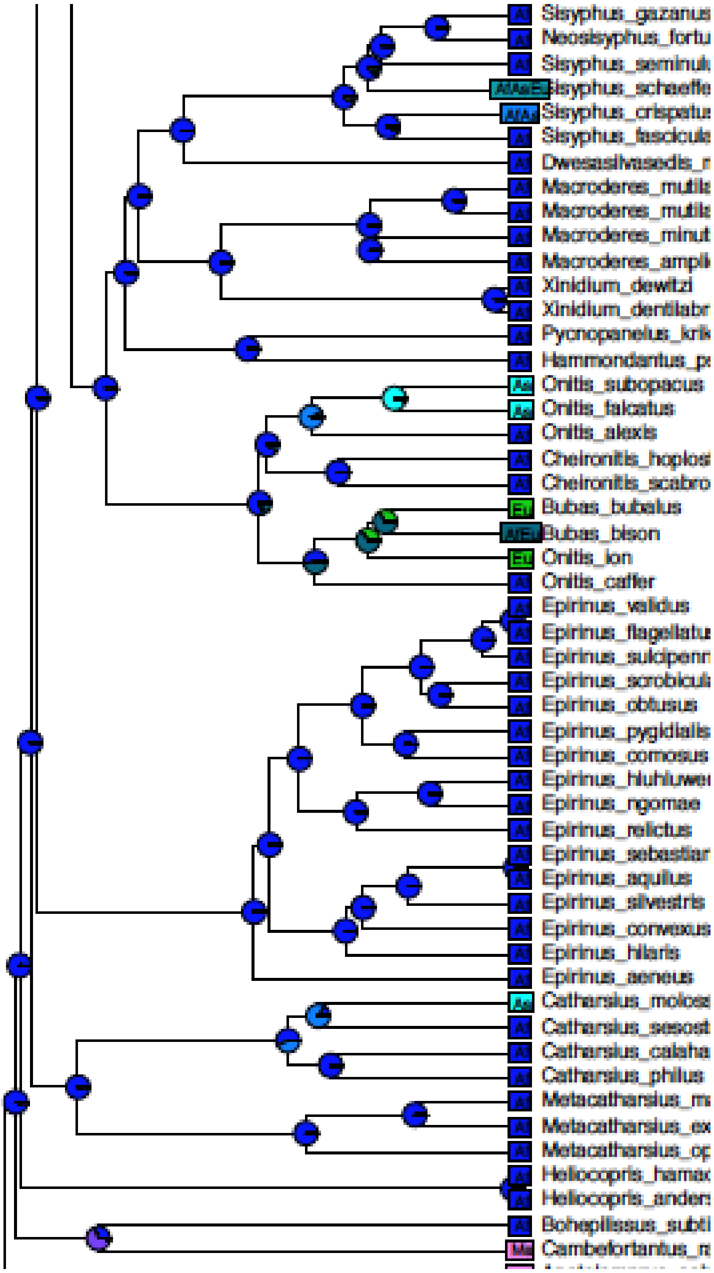


Figure S1 Continued

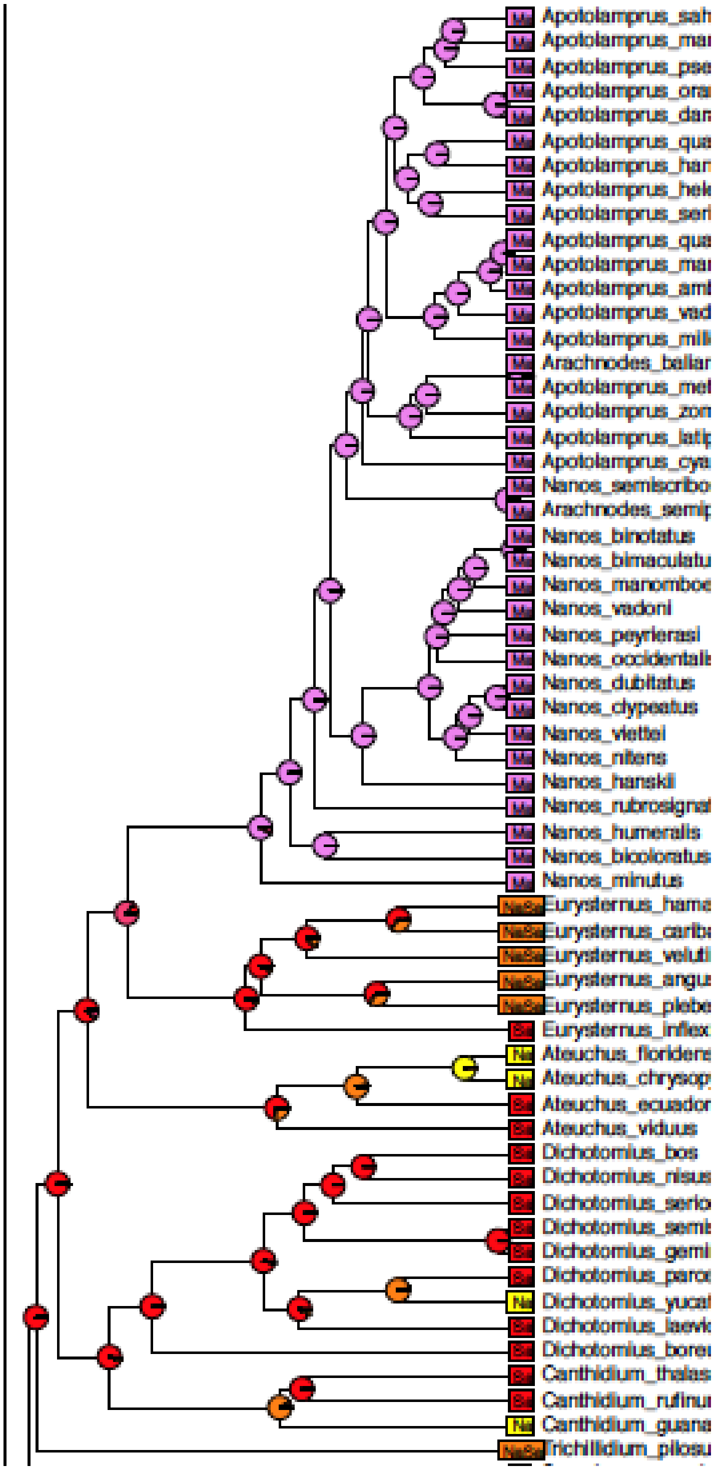


Figure S1 Continued

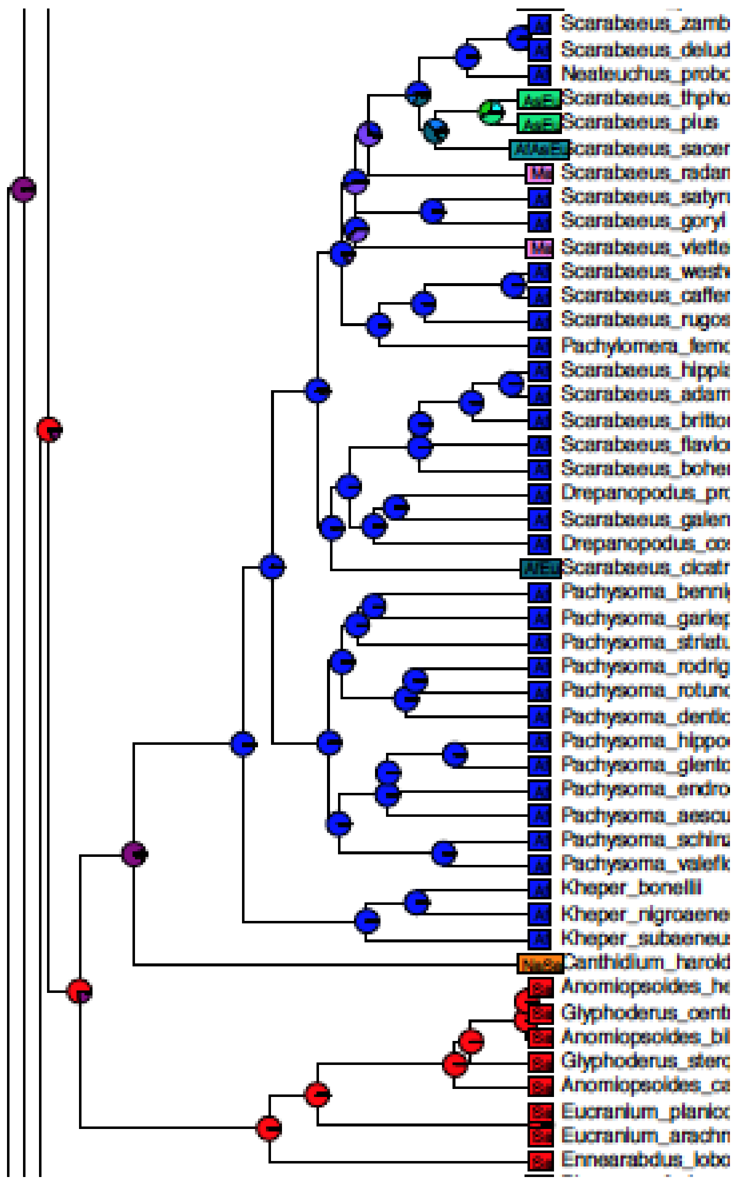


Figure S1 Continued

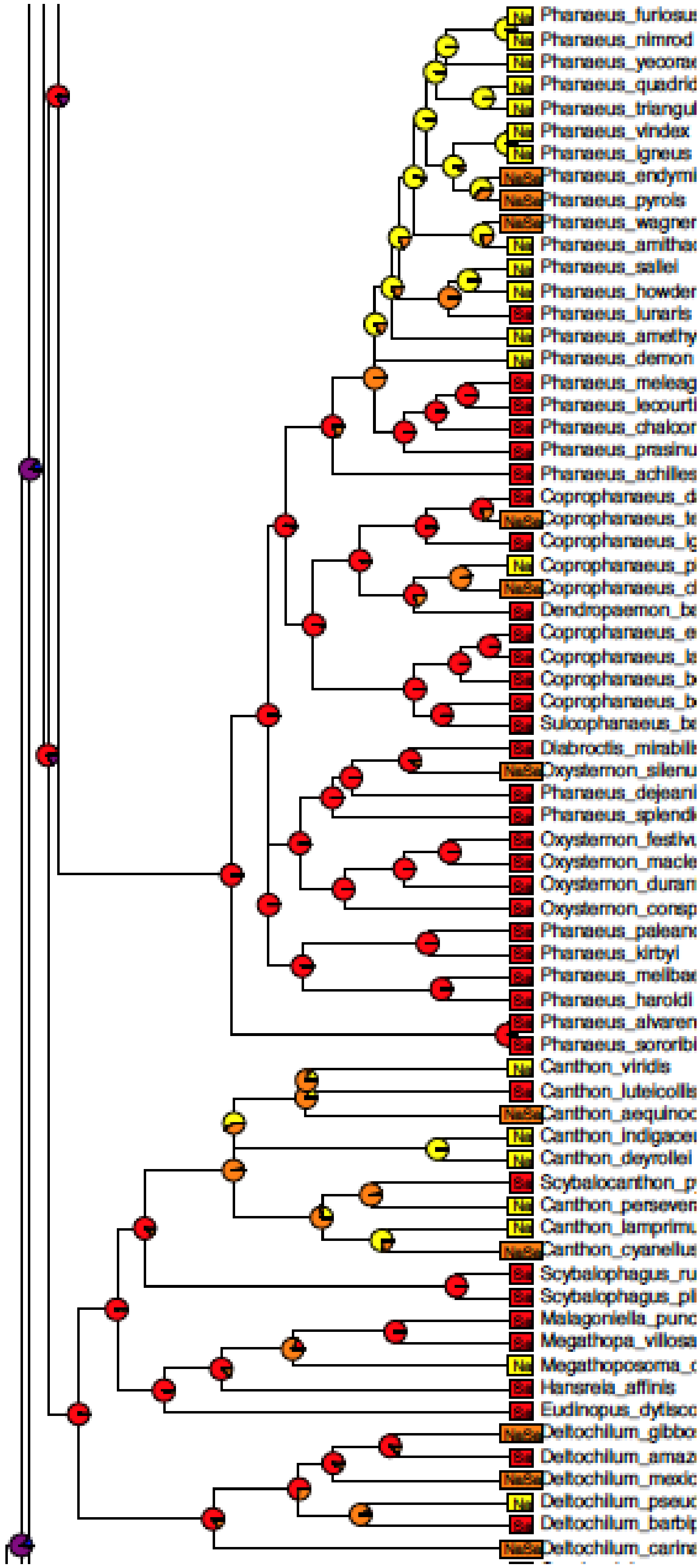


Figure S1 Continued

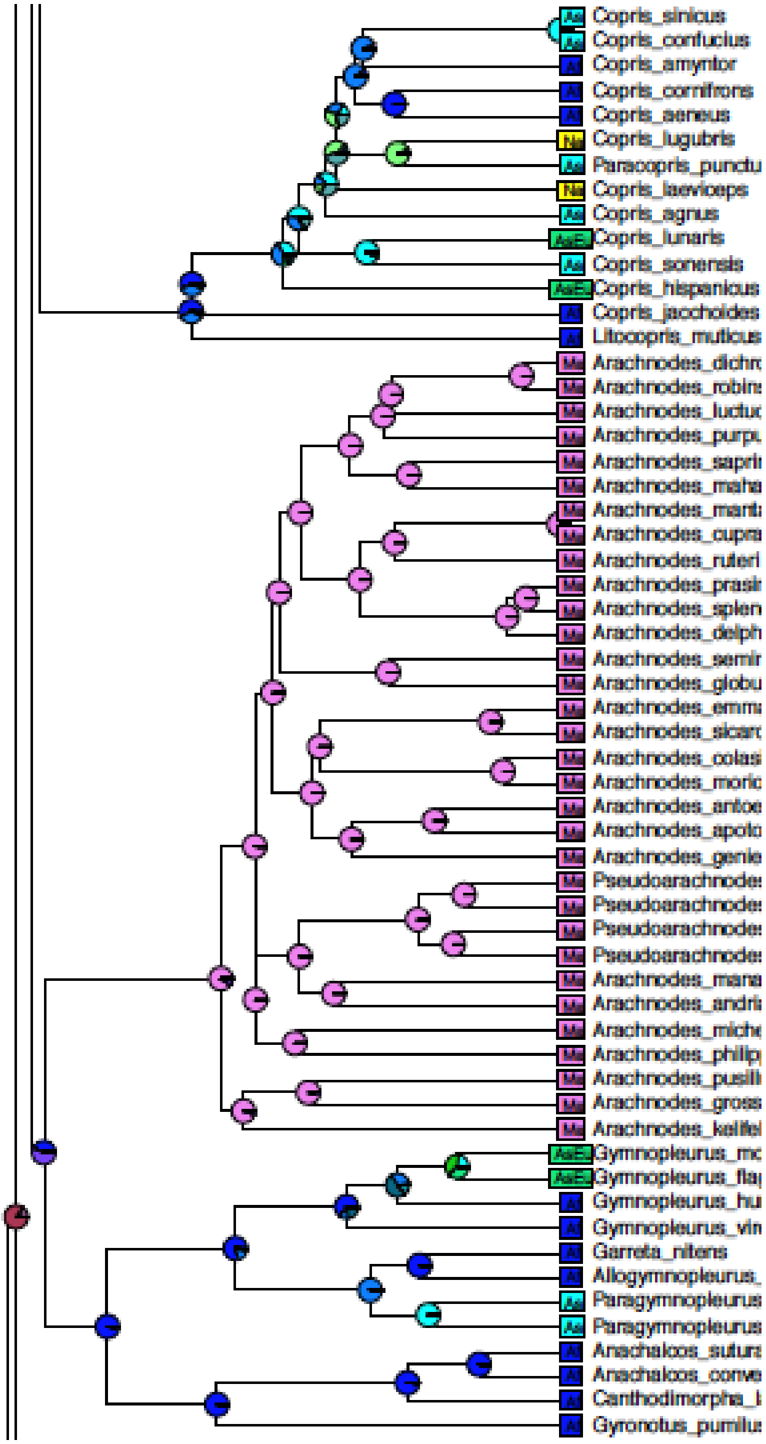


Figure S1 Continued

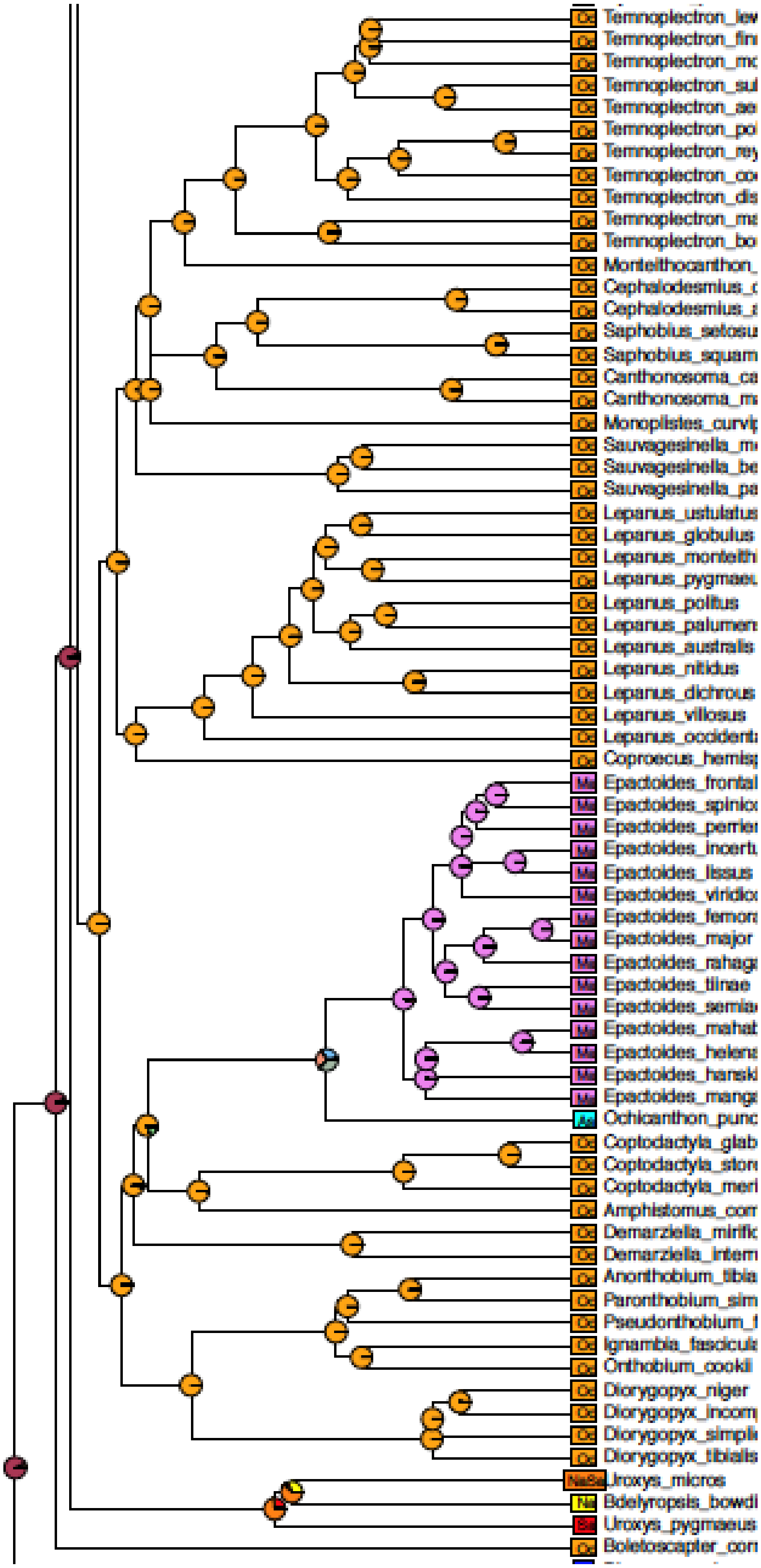


Figure S1 Continued

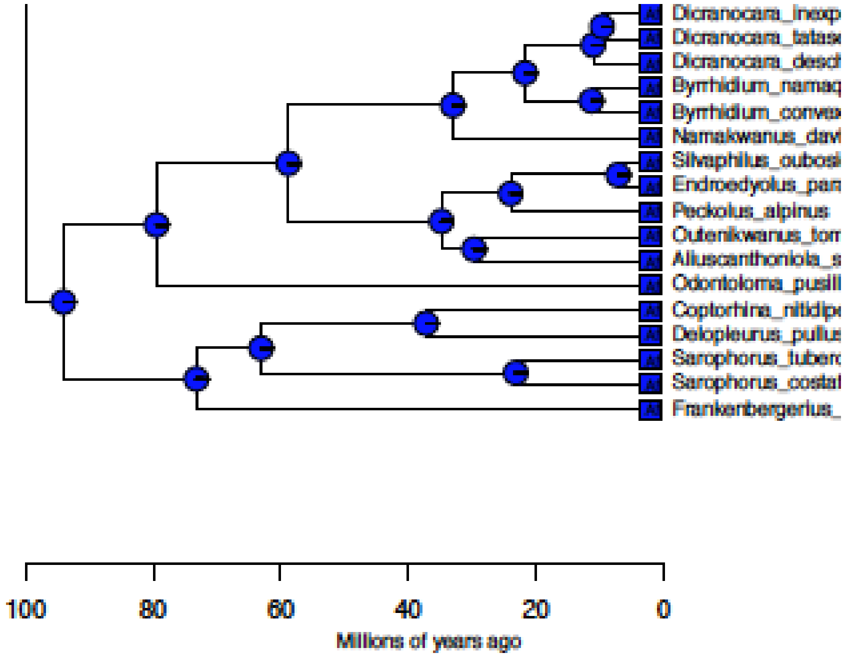


Figure S1 Continued

1
